# Computational modeling establishes mechanotransduction as a potent modulator of the mammalian circadian clock

**DOI:** 10.1101/2023.10.09.561563

**Authors:** Emmet A. Francis, Padmini Rangamani

**Affiliations:** Department of Mechanical and Aerospace Engineering, University of California San Diego, CA 92093, USA

## Abstract

Mechanotransduction, which is the integration of mechanical signals from the cell’s external environment to changes in intracellular signaling, governs many cellular functions. Recent studies have shown that the mechanical state of the cell is also coupled to the cellular circadian clock. To investigate possible interactions between circadian rhythms and cellular mechanotransduction, we have developed a computational model that integrates the two pathways. We postulated that the translocation of the transcriptional regulators YAP/TAZ and MRTF into the nucleus leads to altered expression of circadian proteins. Simulations from our model predict that lower levels of cytoskeletal activity are associated with longer circadian oscillation periods and higher oscillation amplitudes, consistent with recent experimental observations. Furthermore, accumulation of YAP/TAZ and MRTF in the nucleus causes circadian oscillations to decay. These effects hold both at the single-cell level and within a population-level framework. Finally, we investigated the effects of mutations in YAP or lamin A, the latter of which lead to a class of diseases known as laminopathies. Oscillations in circadian proteins are substantially weaker in populations of cells with *in silico* mutations in YAP or lamin A, suggesting that defects in mechanotransduction can disrupt the circadian clock in certain disease states. However, by reducing substrate stiffness, we were able to restore normal oscillatory behavior, suggesting a possible compensatory mechanism. Thus our study identifies that mechanotransduction could be a potent modulatory cue for cellular clocks and this crosstalk can be leveraged to rescue the circadian clock in disease states.

## Introduction

24-hour cycles known as circadian rhythms are a hallmark of life on earth. In mammals, organism-wide circadian oscillations are regulated by signals from the central circadian clock, located in the suprachiasmatic nucleus (SCN) of the hypothalamus (Welsh et al. 2010; Evans 2016). This central clock consists of a population of cells that exhibit oscillations in the expression of circadian proteins, including brain and muscle Arnt-like protein-1 (BMAL1), Period (PER), Cryptochrome (CRY), and nuclear receptor family subfamily 1 group D member 1 (NR1D1), commonly referred to as REV-ERBα (Takahashi 2017; Buhr and Takahashi 2013). These oscillations are synchronized by environmental cues such as light-dark cycles via the process of entrainment. Remarkably, cells from peripheral tissues also exhibit 24-hour patterns of protein expression in these same circadian proteins even when removed from the body, demonstrating the intrinsic nature of the cellular circadian clock. Furthermore, these circadian cycles also influence a wide range of functions in cells; for instance, about 43% of protein-coding genes in mice have been shown to undergo 24-hour cycles in transcription somewhere in the body (Zhang et al. 2014). Recent experimental studies have revealed connections between disease states and disrupted circadian rhythms in different cell types (Shafi and Knudsen 2019; Mason et al. 2020).

Mechanical and chemical cues within a tissue may also serve as circadian clock regulators (Streuli and Meng 2019). While it has long been known that a serum shock synchronizes the circadian oscillations of cells in culture (Balsalobre et al. 1998), it has recently been found that cytoskeletal activity also contributes to clock regulation (Gerber et al. 2013; Esnault et al. 2014). Additional studies have found that the circadian clock is mechanosensitive – the strength of oscillations changes depending on substrate stiffness (Yang et al. 2017; Williams et al. 2018). Xiong et al. recently analyzed this phenomenon in detail, demonstrating that altering the mechanical state of fibroblasts or U2OS cells by either treating with actomyosin inhibitors or seeding cells on substrates of different stiffness leads to distinct changes in the period and amplitude of circadian oscillations (Xiong et al. 2022). In particular, decreases in cytoskeletal activity were generally associated with increases in PER2 oscillation period and amplitude. This was shown to depend on myocardin-related transcription factor (MRTF)-dependent activation of serum response factor (SRF) in the nucleus. In a different study, Abenza et al. found that increased levels of yes-associated protein (YAP) and PDZ-binding motif (TAZ) in the nucleus correlated with significant perturbations to circadian oscillations in fibroblasts (Abenza et al. 2023). This effect was at least partially mediated by transcriptional enhanced associate domain (TEAD). Together, these findings suggest that the cell circadian clock can be affected by the key players in cellular mechanotransduction *viz*. MRTF and YAP/TAZ.

The nuclear translocation of transcriptional regulators such as YAP/TAZ and MRTF is a critical downstream event in cellular mechanotransduction, the process by which cells respond to their mechanical environment via force-dependent changes in biochemical signaling networks. For instance, cells on stiffer substrates show increased levels of focal adhesion kinase (FAK) phosphorylation, leading to downstream changes in cytoskeletal activity and nuclear localization of YAP/TAZ and MRTF. YAP/TAZ and MRTF then bind transcription factors such as TEAD and SRF, causing changes in gene expression. Accordingly, tissue stiffness is a critical determinant of cell behavior, controlling processes from stem cell differentiation to cell migration (Janmey et al. 2020). Additionally, changes to tissue stiffness are often observed in diseases such as cancers, likely contributing to their pathophysiologies (Chin et al. 2016). Here, we examine how changes in tissue stiffness or other mechanical factors might disrupt the circadian clock, perhaps contributing to certain cases of disease progression.

How might cell signaling due to mechanotransduction and circadian oscillations be connected? In this study, we sought to answer this question using computational modeling. Building on the rich history of modeling circadian oscillations in single cells (Asgari-Targhi and Klerman 2019; Leloup and Goldbeter 2003; Lema et al. 2000) and independent models of stiffness-dependent mechanotransduction (Sun et al. 2016; Scott et al. 2021), we developed a model of mechanotransduction-induced perturbations to the cell circadian clock. We constrained our model to recently published experiments, and then used it to investigate the coupling between cell signaling and circadian oscillations in single cells and populations of cells. Finally, we investigated how mutations in YAP or lamin A might impact circadian oscillations. We found that such mutations can significantly weaken circadian oscillations, but this effect can be counteracted by reductions in substrate stiffness. Our model has implications for laminopathies, a class of diseases marked by mutations in the gene for lamin A, as well as diseases marked by changes in local tissue stiffness.

## Results

### Model development

We developed a mathematical model that includes YAP/TAZ and MRTF-mediated mechanotransduction as well as circadian oscillations in the expression of BMAL1, PER/CRY, and REV-ERBα (Figure 1A). The process of mechanosensing leading to nuclear translocation of YAP/TAZ and/or MRTF occurs at a faster timescale compared to changes in the expression of circadian proteins (Figure 1B). Assuming that YAP/TAZ and MRTF levels remain relatively constant over several days, we compute their steady-state nuclear concentrations and use these as inputs to the dynamical model of circadian oscillations. Furthermore, in all cases, we assume that species are well-mixed within a given cellular compartment (plasma membrane, cytosol, nuclear membrane, or interior of the nucleus), allowing us to treat these systems using ordinary differential equations (ODEs) and delay differential equations (DDEs). The mechanotransduction model includes chemical species in all four compartments, whereas the three circadian species are both within the nucleus. We briefly summarize the mechanotransduction and circadian clock modules of our model below.

**Figure 1:**
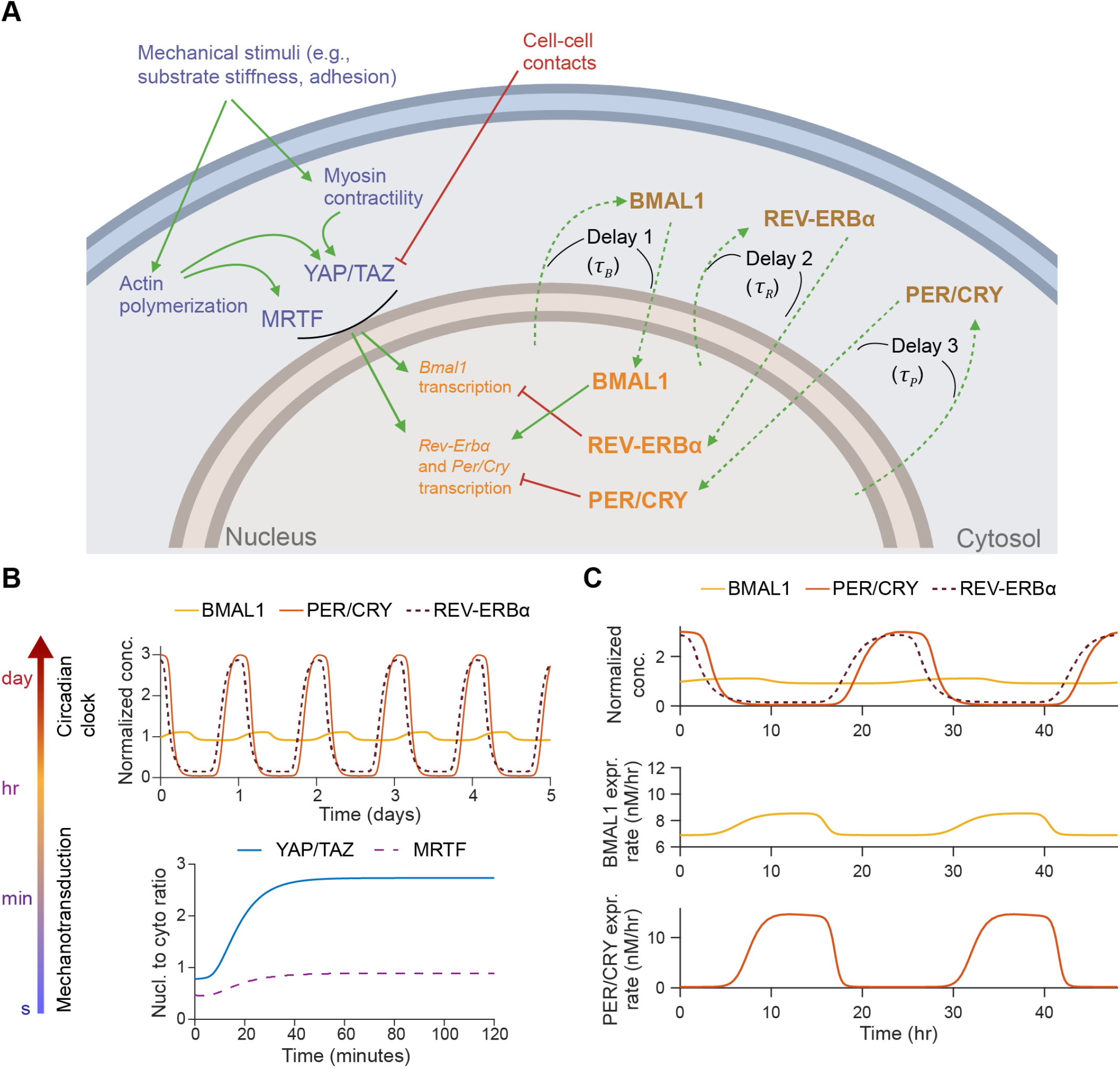
A coupled model of mechanotransduction and the circadian clock. A) Schematic of pathways for YAP/TAZ and MRTF-mediated mechanotransduction and the mammalian circadian clock. Mechanical stimuli such as substrate stiffness and cell-substrate adhesion induce changes in cytoskeletal activity that lead to nuclear translocation of YAP/TAZ and MRTF. Cell-cell contacts also inhibit YAP/TAZ activation via a LATS-dependent pathway. Nuclear YAP/TAZ and MRTF are posited to induce changes in the expression of BMAL1, PER, CRY, and REV-ERBα proteins, altering circadian oscillations. In parallel, increases in nuclear BMAL1 enhance transcription of *Per, Cry*, and *Rev-ERb*α, while increases in nuclear REV-ERBα inhibit transcription of *Bmal1* and increases in nuclear PER/CRY inhibit transcription of *Per, Cry*, and *Rev-Erb*α. Dashed lines indicate delays associated with transcription, translation, post-translational modifications, and nuclear translocation of BMAL1, PER/CRY, and REV-ERBα. The schematic was created with Biorender.com. B) Separation of timescales from mechanotransduction pathways to circadian oscillations. Events downstream of mechanical stimuli lead to changes in YAP/TAZ and MRTF nuclear concentrations within tens of minutes (lower plot), whereas changes in the concentration of circadian proteins are extended over hours to days (upper plot). C) Simulated dynamics of single-cell circadian oscillations. BMAL1, PER/CRY, and REV-ERBα all oscillate with periods close to 1 day. Rates of expression of BMAL1 and PER/CRY are depicted in lower plots.

### YAP/TAZ and MRTF mechanotransduction model

Our mechanotransduction model is built from the model originally developed by Sun et al. (Sun et al. 2016) and then expanded by Scott et al. (Scott et al. 2021). In brief, cell-substrate adhesion leads to substrate-stiffness dependent phosphorylation of focal adhesion kinase (FAK) adjacent to the cell membrane, triggering downstream activation of the GTPase RhoA. RhoA then activates both Rho kinase (ROCK) and the formin mDia, which promote myosin activity and actin polymerization, respectively. ROCK also indirectly promotes actin polymerization by activating LIM kinase (LIMK), which inactivates cofilin, an F-actin-severing protein. Upon activation, myosin and actin form stress fibers, leading to the dephosphorylation of YAP and TAZ in the cytosol and allowing their translocation into the nucleus. When simulating different cell densities, we also assume that cell-cell contacts induce increased phosphorylation of YAP/TAZ via a LATS-dependent pathway, causing more YAP/TAZ to remain cytosolic. Actin polymerization also permits the release of G-actin-sequestered MRTF, which can then translocate into the nucleus. As in Scott et al. (Scott et al. 2021), we assume that the opening of nuclear pore complexes (NPCs) depends on the phosphorylation state of lamin A; in particular, lamin A is increasingly dephosphorylated on stiffer substrates, leading to its incorporation into the nuclear lamina and concomitant stretching of NPCs (Swift et al. 2013). This allows for increased transport of both YAP/TAZ and MRTF into the nucleus. Scott et al. did not include MRTF (Scott et al. 2021), whereas Sun et al. did not consider contributions of lamin A and the NPC to the nuclear transport of MRTF and YAP/TAZ (Sun et al. 2016). Here, we combine elements of both models to fully examine both YAP/TAZ and MRTF nuclear translocation. This mechanotransduction model includes 25 species, which can be reduced to 13 ordinary differential equations by accounting for mass conservation. We solve for steady state values by setting each differential equation equal to zero; the resulting expressions for each steady-state quantity are given in Table S2. For all YAP/TAZ-related parameters, we use the values previously estimated for mammalian cells (Scott et al. 2021) (see Table S3), and MRTF-specific parameters and sensitivities to different treatment conditions were calibrated as described below. Baseline values for MRTF rate constants were set to match the values used for YAP/TAZ where applicable (Table S4).

### Circadian clock model

There is a vast body of literature on modeling cell circadian oscillations (reviewed in Asgari-Targhi and Klerman 2019). Here, rather than attempt to recapitulate the mammalian circadian clock in full mechanistic detail, we develop a reduced-order model, including the dynamics of positive and negative clock regulators through three representative species, *B* (nuclear concentration of BMAL1), *P* (nuclear concentration of PER and CRY proteins), and *R* (nuclear concentration of REV-ERBα). This reduced approach is made possible by using DDEs, as in multiple previous circadian models of varying complexity (Lema et al. 2000; Sriram et al. 2006; Smolen et al. 2001). In addition to these DDE models, we integrate knowledge from previous ODE-based models of the mammalian circadian clock (Leloup and Goldbeter 2003; Mirsky et al. 2009; Forger and Peskin 2003).

In our model, BMAL1 induces the expression of REV-ERBα and PER/CRY after translocating into the nucleus and forming a complex with CLOCK (circadian locomotor output cycles kaput). CLOCK levels remain relatively constant over time (Shearman et al. 2000), and so it is not treated as a separate dynamical variable here. Upon its expression and transport into the nucleus, REV-ERBα inhibits the expression of BMAL1 (Takahashi 2017). This effect is modeled as an inhibition with delay *τ*_*B*_, which represents the time between changes in transcription and associated changes in the nuclear concentration of BMAL1. This includes the time for transcription, translation, post-translational modifications, and shuttling of BMAL1 into the nucleus. As a starting estimate for this parameter, we use the time gap between peak BMAL1 transcription and peak nuclear BMAL1, which has been measured at 12-16 hours in mouse fibroblasts (Tamaru et al. 2003). Writing *R*[*t* − *τ*_*B*_] as the nuclear concentration of REV-ERBα at time *t* − *τ*_*B*_, the REV-ERBα-regulated BMAL1 expression level is:

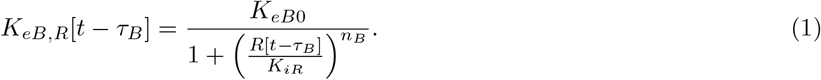

Similarly, writing the delay between changes in PER/CRY (*P*) transcription and eventual changes in nuclear PER/CRY as *τ*_*P*_ (approximated to be 6-9 hours based on measurements in mouse cells (Yagita et al. 2001; Yagita et al. 2002; Lee et al. 2001)), the BMAL1-regulated expression of PER/CRY is:

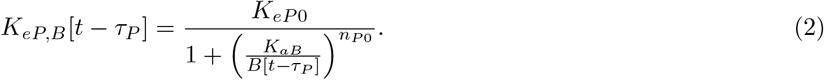

PER/CRY, in turn, binds to the BMAL1-CLOCK complex in the nucleus, inhibiting its upregulation of genes controlled by E-box enhancer elements (including both *Per/Cry* and *Rev-Erb*α). We write PER/CRY self-inhibition as:

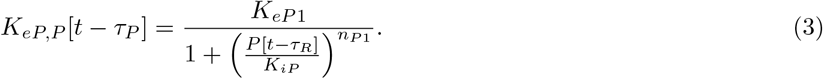

For simplicity, we model changes in REV-ERBα expression in the same manner as PER/CRY. Because both genes are regulated by E-box enhancers, we assume the expression terms are proportional to one another:

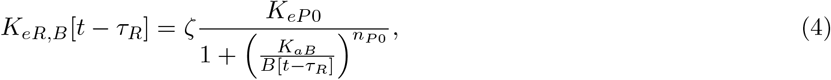

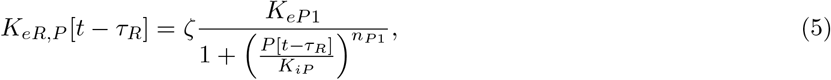

where *ζ* is a proportionality constant and *τ*_*R*_ is the delay between REV-ERBα transcription and changes in nuclear REV-ERBα.

Furthermore, we assume that nuclear YAP/TAZ ([*Y*_*nuc*_]) and MRTF ([*M*_*nuc*_]) independently regulate the expression of circadian proteins via TEAD and SRF, respectively. In reality, these processes may not be fully independent of the nuclear concentrations of BMAL1, PER/CRY, and REV-ERBα, but sufficient mechanistic details are not available to justify a particular functional dependence. These YAP/TAZ and MRTF-dependent expression terms, *K*_*eB*2_, *K*_*eP* 2_, and *K*_*eR*2_ are written as sums of Hill equations:

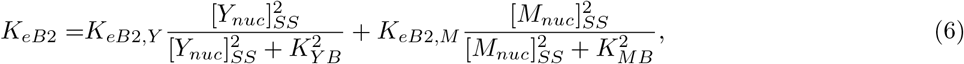

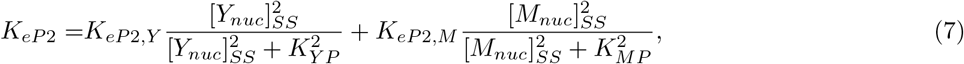

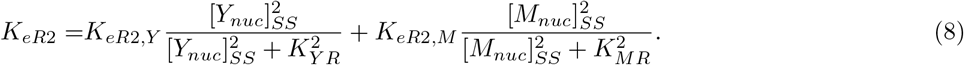

For each of these expression functions in Equations (6) to (8), the Hill coefficient was fixed at 2 as it was not found to affect the qualitative behavior of our model (Figure S1). It should be noted that [*Y*_*nuc*_]_*SS*_ and [*M*_*nuc*_]_*SS*_ in Equations (6) to (8) are steady-state values; it is implicitly assumed that the mechanotransduction signaling pathways have reached steady state prior to the start of our simulations.

We write out the full set of DDEs, including decay terms for each species individually:

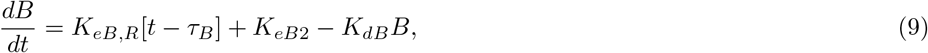

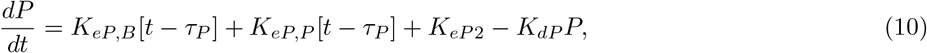

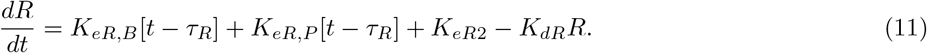

For a direct comparison of our model to experimental measurements of luminescence from cells expressing a PER2 promoter-driven luciferase reporter (PER2::Luc) (Yoo et al. 2004; Xiong et al. 2022), we explicitly modeled luciferase dynamics. Assuming the expression of luciferase scales directly with PER2 expression and cytosolic luciferase decays with rate *K*_*dL*_ (measured in Feeney et al. 2016), its dynamics are given by integrating the following expression:

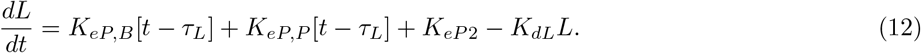

Altogether, this Circadian clock model involves 28 free parameters as summarized in Table S1. Baseline values were chosen to match the single DDE system in Lema et al. (Lema et al. 2000) or experimentally measured delays (Yagita et al. 2001; Yagita et al. 2002; Lee et al. 2001; Tamaru et al. 2003), or they were empirically set from early testing of the model.

### Model calibration

We assessed model sensitivity to each parameter by computing the total-order Sobol’ indices associated with the luciferase reporter’s oscillation period and amplitude (Figure S2). Although the amplitude of the reporter was mostly sensitive to a few parameters (Figure S2B), oscillation period was sensitive to all free parameters in our model (Figure S2A). Accordingly, we estimated all free parameters, including circadian parameters, MRTF transport parameters, and inhibitor treatment-related parameters, rather than fix any at arbitrary default values. YAP/TAZ-related parameters were fixed to the values previously calibrated to experimental data (Scott et al. 2021). Given the inherent variability in biological systems, we estimated probability distributions rather than single values for all parameters in Tables S4 and S1 using Bayesian parameter estimation (Linden et al. 2022). We utilized measurements of PER2::Luciferase mouse primary fibroblasts in Xiong et al. to calibrate our model (Xiong et al. 2022, Table S6). We minimized the error between model predictions and experimental measurements for circadian oscillations after changing substrate stiffness (Figure 2A) or treating cells with a ROCK inhibitor (Y27632) (Figure 2B), cytochalasin D (Figure 2C), latrunculin B (Figure 2D), or jasplakinolide (Figure 2E). These treatments introduce distinct effects on the cytoskeleton or associated signaling pathways while only introducing five additional free parameters. Overall, simulations using parameters sampled from their posterior distributions (Figure S2C) provided a reasonable fit to experimental data (Figure 2A-E). More details on sensitivity analysis and parameter estimation are given in Methods (Section “Sensitivity analysis and parameter estimation“).

**Figure 2:**
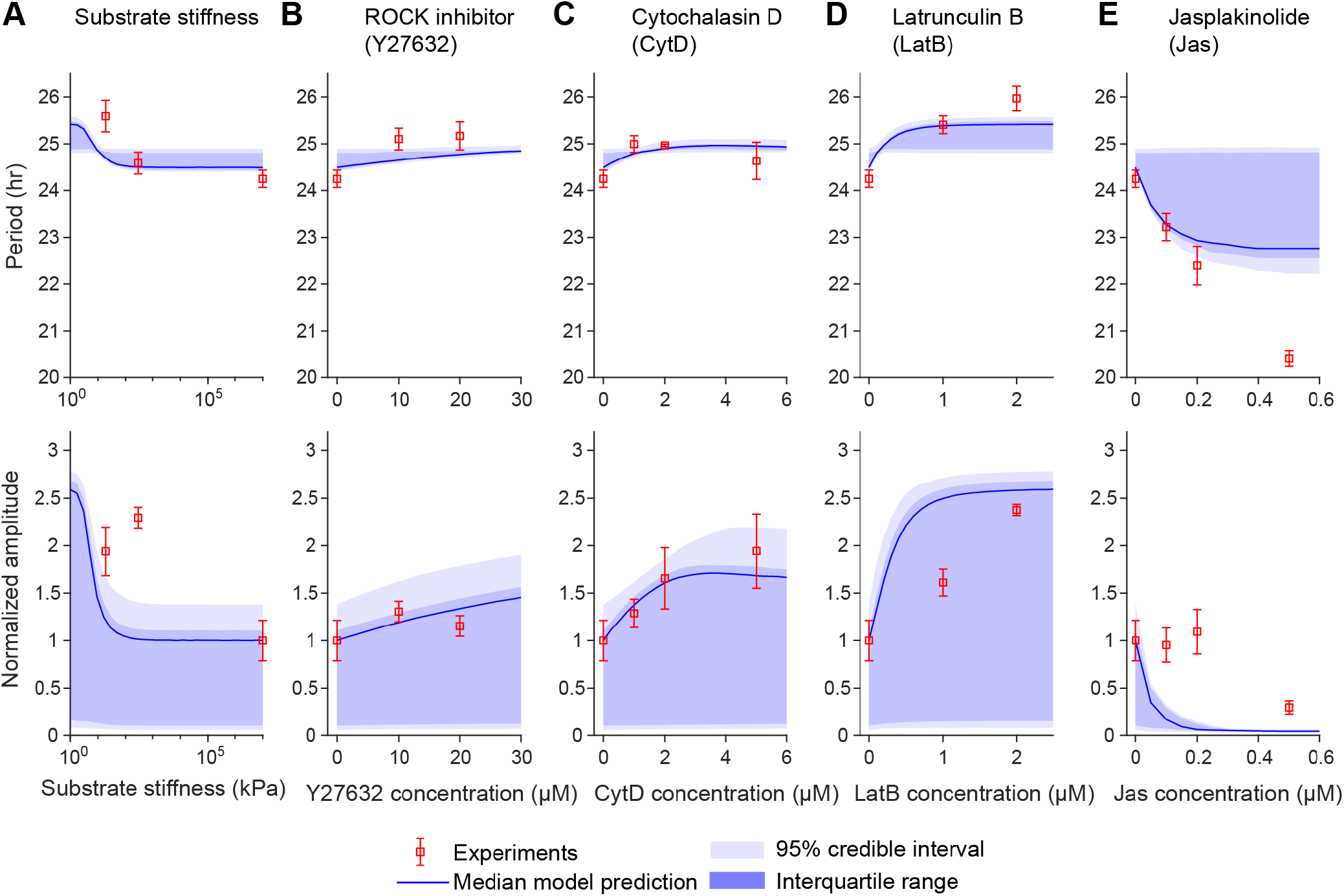
Bayesian parameter estimation for mechanotransduction-circadian model. A-E) Predicted oscillation period and amplitude in 5 experiments from Bayesian parameter estimation. Experimental data from Xiong et al. (Xiong et al. 2022) and model values are plotted for the effects of substrate stiffness (A), ROCK inhibitor (Y27632) treatment (B), cytochalasin D treatment (C), latrunculin B treatment (D), and jasplakinolide treatment (E). Experimental error bars denote standard deviation. For the model, the darker blue region indicates interquartile range and the lighter blue region spans the 95% credible interval (region from the 2.5 percentile to 97.5 percentile), based on sampling from the posterior distributions for parameter values. 1000 samples were generated to compute the estimates in this figure. In all cases, amplitude is normalized to that associated with the control condition (untreated cells on glass).

### Mechanical factors alter the stability and dynamics of circadian oscillations

We examined the behavior of our model using the set of maximum *a posteriori* parameters from Bayesian parameter estimation (Tables S1 and S4). This corresponds to a single representative cell that exhibits behavior close to the mean population measurements from Xiong et al. 2022.

We first tested the effects of changing substrate stiffness. F-actin, cytosolic stiffness, nuclear YAP/TAZ, and nuclear MRTF all increase consistently as a function of substrate stiffness (Figure 3A). In turn, we observed marked changes to circadian oscillations (Figure 3B); as substrate stiffness increases, the period and amplitude of circadian oscillations also increase, but the overall oscillatory nature remains consistent. These findings agree well with the data used for model calibration (Xiong et al. 2022) and validate the coupling between our mechanotransduction module and the circadian module.

**Figure 3:**
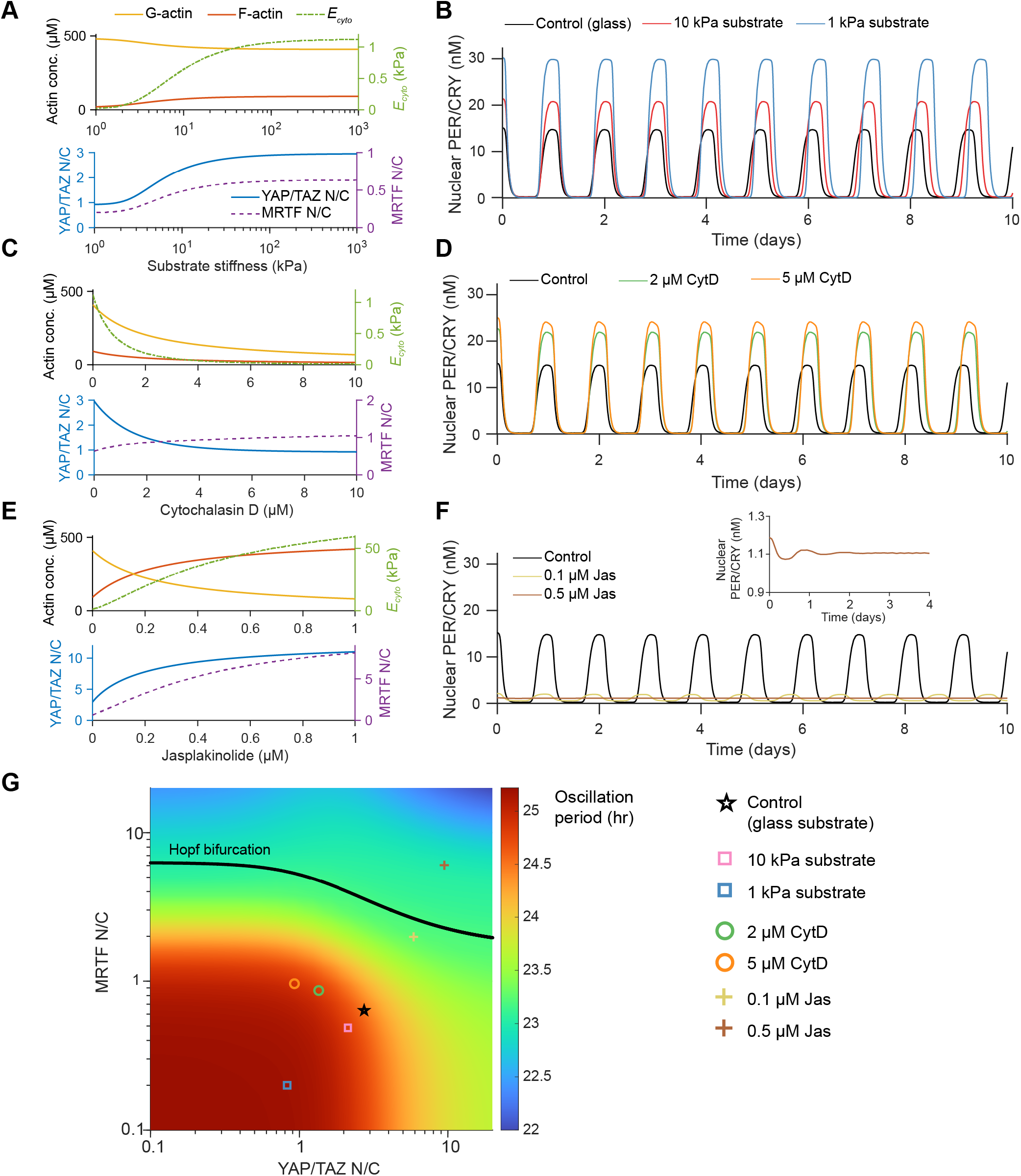
Changes to circadian oscillations due to altered cytoskeletal activity. A) Substrate-stiffness-dependent changes in actin, cytosolic stiffness (*E*_*cyto*_), nuclear YAP/TAZ, and nuclear MRTF. B) Substrate-stiffness-dependent changes to oscillations in nuclear PER/CRY. C) Cytochalasin-D-induced changes to actin, cytosolic stiffness, and nuclear levels of YAP/TAZ and MRTF. D) Cytochalasin-D-induced changes to oscillations in nuclear PER/CRY. E) Jasplakinolide-induced changes to actin, cytosolic stiffness, and nuclear levels of YAP/TAZ and MRTF. F-actin increases dramatically upon jasplakinolide treatment, leading to substantial nuclear accumulation of YAP/TAZ and MRTF. F) Jasplakinolide-induced changes to oscillations in nuclear PER/CRY. Increased actin polymerization leads to a decrease in the oscillation period along with a decay in circadian oscillations (inset). G) Bifurcation diagram showing the location of the Hopf bifurcation and the dependence of the circadian oscillation period on nuclear YAP/TAZ and MRTF. All points below the Hopf bifurcation curve correspond to sustained oscillations, whereas points below exhibit decaying oscillations. Individual markers plotted in the YAP/TAZ-MRTF phase plane correspond to the test cases shown in panels A-F.

Next, we investigated how two different cytoskeleton-targeting drugs, cytochalasin D and jasplakinolide, affect circadian oscillations. Cytochalasin D acts by capping existing actin filaments and inducing dimerization of G-actin. We modeled this in a similar manner to Wakatsuki et al. 2001, as fully described in Methods and Table S5. Because cytochalasin D treatment decreases the concentration of both F-actin and G-actin, it induces opposite effects on nuclear YAP/TAZ and MRTF (Figure 3C). Specifically, nuclear YAP/TAZ decreases whereas nuclear MRTF increases due to the G-actin dimerization, in agreement with experiments (Finch-Edmondson and Sudol 2016; Miralles et al. 2003). Jasplakinolide stabilizes actin filaments and also induces polymerization (Bubb et al. 2000). We modeled this by assuming that the polymerization rate increases and the depolymerization rate decreases as a function of jasplakinolide concentration (Section S3.3, Table S5). As a result, F-actin accumulates in the cytosol, leading to increased levels of YAP/TAZ and MRTF in the nucleus (Figure 3E).

As expected, these inhibitor treatments have distinct effects on circadian oscillations. Similar to the effects of lower substrate stiffness, cytoskeletal inhibition by cytochalasin D leads to increased circadian oscillation period and amplitude (Figure 3D). Conversely, when actin polymerization is enhanced via jasplakinolide treatment, YAP/TAZ and MRTF accumulate in the nucleus, leading to a slight decrease in oscillation period and rapidly decaying oscillations (Figure 3F). This suggests that circadian oscillations are robust at lower levels of nuclear YAP/TAZ and MRTF, but upon accumulation of these factors in the nucleus, oscillations can decay over time. The location of this transition from stable amplitude to damped oscillations (the Hopf bifurcation) in the YAP/TAZ-MRTF plane is shown in Figure 3G.

This general behavior, in which a Hopf bifurcation occurs for higher nuclear concentrations of YAP/TAZ and MRTF, is not exclusive to the single set of parameters used here. This feature of the model is conserved for wide range of *K*_*dB*_, *K*_*dP*_, and *K*_*dR*_ tested in Figure S3. In fact, even a further reduced version of our model (examined analytically in Section S5) shows this same trend: sustained circadian oscillations only occur when mechanically induced expression remains below a certain threshold.

### Model captures population-level variability in circadian oscillations

We next sought to capture population-level variability in circadian oscillations in our model. To incorporate population-level features, we focused on cell-to-cell variation of kinetic parameters. Taking inspiration from computational studies in cardiomyocytes (Ni et al. 2018; Kernik et al. 2019) and stochastic models of genetic oscillators (Veliz-Cuba et al. 2015), we represented this natural variability in kinetic parameters by generating a model cell population in which individual cells have kinetic parameters drawn from probability distributions. These distributions were directly adopted from the posterior distributions derived from Bayesian parameter estimation or, in the case of fixed parameters in the YAP/TAZ model, were assumed to be log-normal (Section “Simulation of cell populations“, Figure S2C). Furthermore, to capture additional sources of measurement noise, we added Gaussian white noise to each time series after solving the DDEs.

We first verified that simulations using this modified model still captured the overall trends seen at the single-cell level (Figure 3) while displaying increased variability. In this case, we analyze oscillations in REV-ERBα rather than PER/CRY for a direct comparison with the population-level data from Abenza et al. (Abenza et al. 2023). Plotting population REV-ERBα dynamics as a kymograph, we observe that circadian oscillations appear to be stronger on a soft substrate compared to cells on glass (Figure 4A-B), similar to the decreased oscillation amplitude observed on stiff substrates in the single-cell case (Figure 3A). In further agreement with single-cell observations, cells on stiffer substrates exhibit decreased oscillation period on average (Figure 4C).

**Figure 4:**
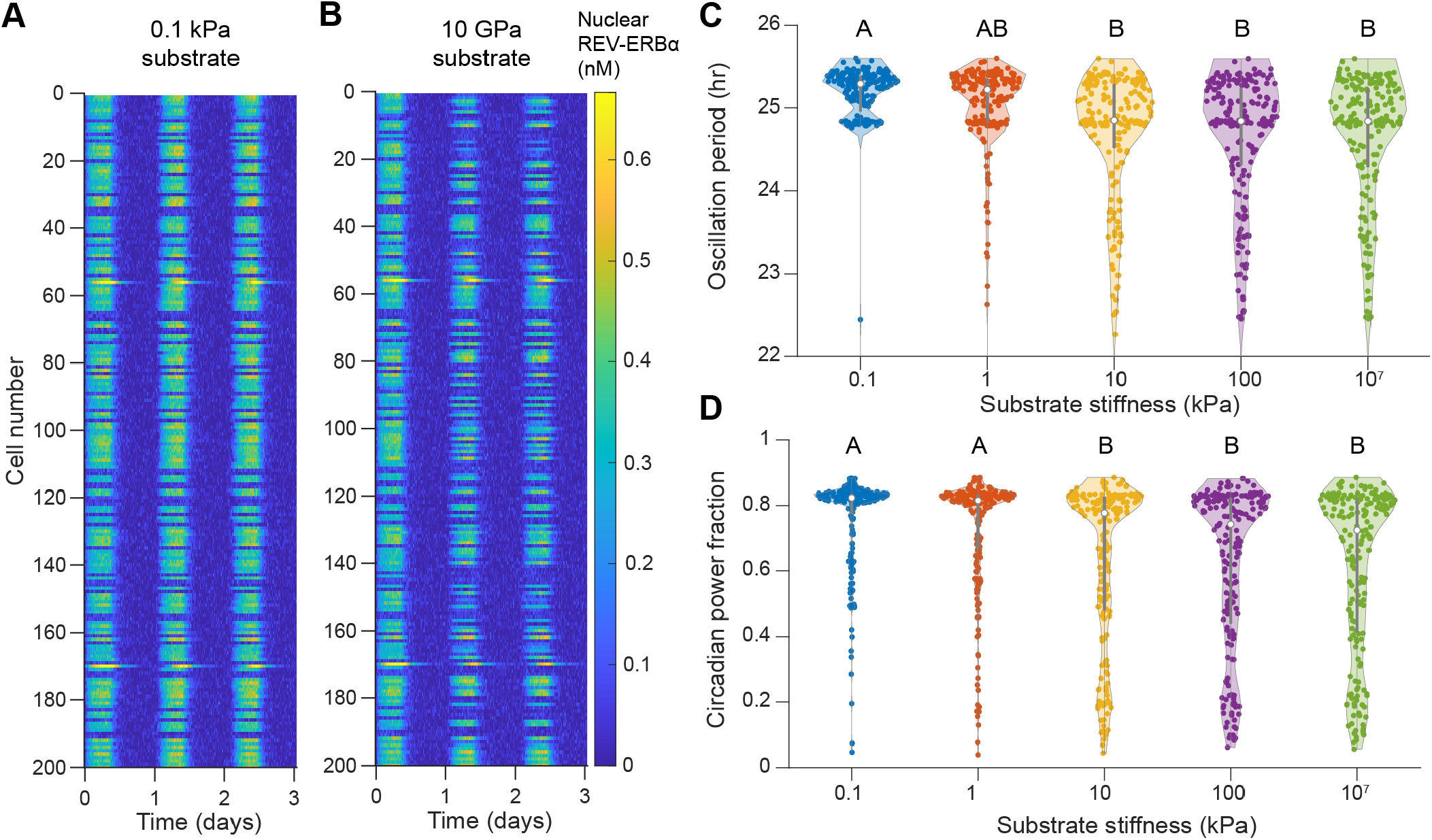
Behavior of circadian oscillations in model cell populations with variability. A) Kymograph depicting oscillations in nuclear REV-ERBα for 200 model cells on a soft (0.1 kPa) substrate. Cells exhibit consistent circadian oscillations, while variability is observed at the level of the population. B) Kymograph depicting oscillations in nuclear REV-ERBα for 200 model cells on glass (10 GPa). Most cells show consistent, lower magnitude oscillations; in some cases, oscillations are weaker or nonexistent. C-D) Distribution of oscillation period (C) and circadian power fraction (D) for cell populations on different substrate stiffnesses. Central points and error bars in violin plots denote the median and interquartile range. Compact letter display is used to denote statistical significance, where groups sharing a letter are statistically similar according to ANOVA followed by Tukey’s *post hoc* test with a significance threshold of *p* = 0.05. Violin plots were generated using Violinplot in MATLAB (Bechtold 2016)

To quantify disruptions to regular circadian oscillations, we used a metric called the circadian power fraction (originally defined in Abenza et al. 2023), which ranges from 0 (no circadian oscillations) to 1 (perfect sinusoidal circadian oscillations). This metric corresponds to the fraction of the circadian power spectrum contained within a close window of the population-level average frequency, as detailed in Methods (Section “Calculation of the circadian power frac-tion“). Lower substrate stiffnesses are generally associated with higher circadian power fractions, whereas increased accumulation of YAP/TAZ and MRTF in the nucleus on stiff substrates results in lower circadian power fractions (Figure 4D). These lower circadian power fractions indicate significantly disrupted oscillations, which manifest as weaker oscillations that are drowned out by Gaussian noise in our simulations (Figure 4B). We note that Abenza et al. did not observe the large increase in circadian power fraction on softer substrates that we report here (Abenza et al. 2023); however, the average YAP/TAZ nuclear to cytosolic ratio also appears to remain elevated for cells on soft substrates in their study, perhaps indicating their cells remained in a mechanically active state not accounted for in our model. Alternatively, other factors could contribute to changes in the expression of circadian proteins in this case; this option is further explored below.

### Nuclear abundance of YAP/TAZ and MRTF correlate with the strength of circadian oscillations across treatment conditions

Next, we considered how other changes in the mechanical environment or distinct pharmacological treatments that disrupt the cytoskeleton can impact the circadian power fraction. We tested the set of conditions examined experimentally in Abenza et al. 2023, including changes in substrate stiffness, cell density, or cell adhesion area, and treatment with cytochalasin D, latrunculin A, or blebbistatin. While several of these treatments were considered in our model calibration and initial testing, here we directly leverage the model against this separate set of experiments and we mainly focus on the strength of oscillations at the cell population level. The assumed effects of each treatment are given in Table S5, with further details in Methods. In brief, increased cell density and restriction of cell adhesion area (1600 or 900 µm^2^ micropatterns) were both assumed to reduce the amount of FAK phosphorylation due to less cell-substrate contact (Section S3.4). Increased cell density was also assumed to increase the YAP/TAZ phosphorylation rate due to activation of LATS (Table S5). Latrunculin A and blebbistatin treatment decreased the F-actin polymerization rate and the formation of active stress fibers, respectively (Section S3.2, Table S5).

As expected, all of these treatments decrease nuclear YAP/TAZ and MRTF on average, with the exception of cytochalasin D, which increases the nuclear concentration of MRTF as described above (Figure 5). Given our previous results, we expect any treatments decreasing nuclear YAP/TAZ and/or MRTF to enhance circadian oscillations. Indeed, in all cases except cytochalasin D treatment, we do observe an increase in the median circadian power fraction (Figure 5).

**Figure 5:**
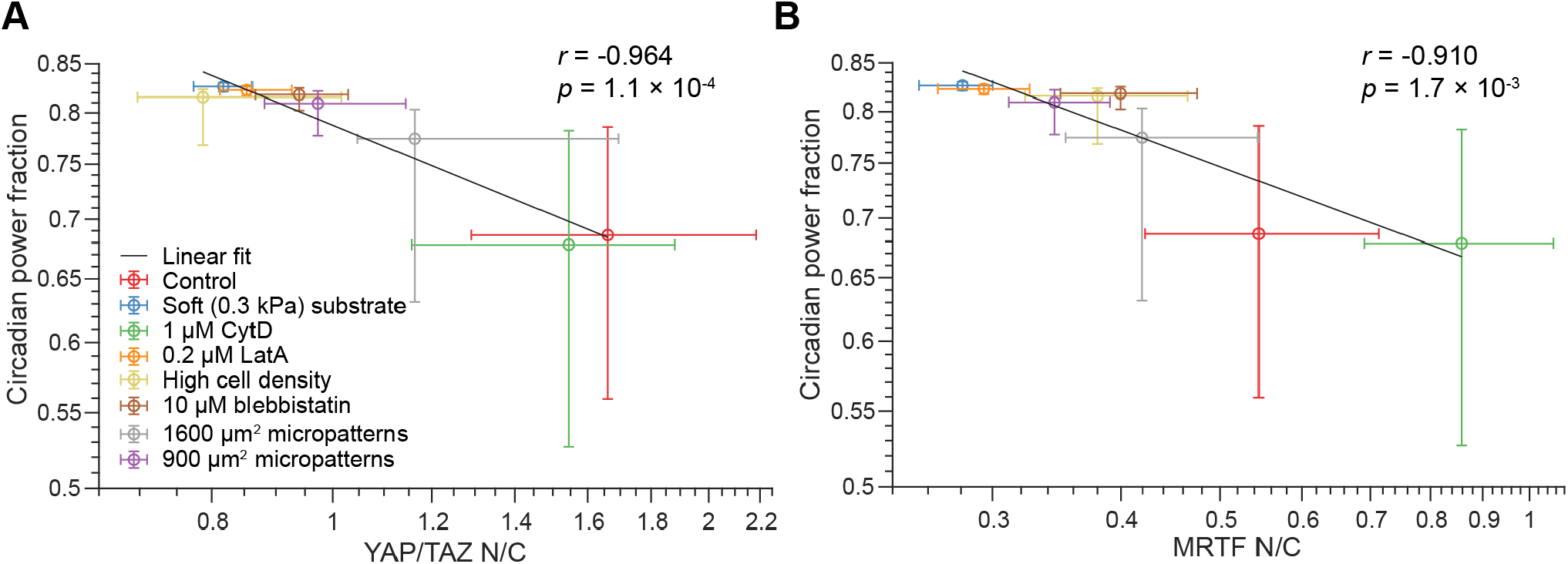
Correlation between circadian power fraction and nuclear to cytosolic ratios (N/C) of YAP/TAZ or MRTF. Populations of cells were subjected to the treatments from Abenza et al. 2023 and circadian power fraction was plotted against YAP/TAZ N/C (A) and MRTF N/C (B). Data points denote median values and error bars denote the range from the 40th to the 60th percentile for 500 model cells. In each case, Pearson’s correlation coefficient (*r*) was assessed for the correlation between median power fraction and median N/C on the log scale. *r* and the *p*-value associated with the null hypothesis, *r* = 0, are included on each graph. To generate distinguishable colors for each condition, we used the linspecer tool in MATLAB (Lansey 2023).

In Abenza et al. 2023, the authors measured a significant correlation between YAP/TAZ nuclear to cytosolic ratio (N/C) and the circadian power fraction, but no such correlation for MRTF N/C. In agreement with their findings, we see a strong correlation between the power fraction and nuclear YAP/TAZ (Figure 5A). However, we also see a significant correlation of the power fraction with nuclear MRTF (Figure 5B). Notably, this correlation is weaker than that with YAP/TAZ (see correlation coefficient statistics in Figure 5). Additionally, we noted that cells exhibited low circadian power fractions even on soft substrates in Abenza et al. 2023. Accordingly, we postulated that an additional mechanism might disrupt circadian oscillations on soft substrates. We tested this by increasing the baseline expression of BMAL1 and PER/CRY in cells on a soft substrate and observed that the correlation with MRTF nuclear abundance was no longer significant, whereas the correlation with nuclear YAP/TAZ remains significant (Figure S4). This higher sensitivity of circadian oscillation amplitude to changes in YAP/TAZ is consistent with the fact that, among mechanotransduction-circadian coupling parameters, circadian oscillation amplitude is most sensitive to *K*_*eP* 2,0_ (Figure S2B). Furthermore, we can clearly see that oscillation amplitude changes more quickly with YAP/TAZ over the YAP/TAZ-MRTF phase plane over the region corresponding to the conditions tested here (Figure S3B).

### Computational predictions establish that YAP or LMNA mutations can significantly disrupt circadian oscillations

Both our single-cell model and our population-level models predict that YAP/TAZ and MRTF nuclear concentrations modulate the strength and stability of circadian oscillations (Figure 3G, Figure 4, Figure 5). We therefore reasoned that mutations affecting the associated pathways for nucleo-cytoplasmic transport could strongly impact circadian oscillations. We first tested this by considering the overexpression of the YAP mutant 5SA-YAP; such overexpression in fibroblasts was previously found to result in a large reduction in the circadian power fraction (Abenza et al. 2023). This mutant has several phosphorylation sites removed, causing it to accumulate at abnormal levels in the nucleus. We tested this condition in our model by introducing a new species, 5SA-YAP, which behaved the same as wild-type YAP/TAZ, but had a phosphorylation rate equal to zero (Table S5). In these tests, the amount of wild-type YAP/TAZ and mutant YAP were each equal to the amount of YAP/TAZ in control cells, resulting in twice the overall YAP/TAZ per cell. Due to the combined effects of overexpression and decreased phosphorylation, YAP/TAZ nuclear concentration significantly increases compared to wild type cells (Figure 6A).

**Figure 6:**
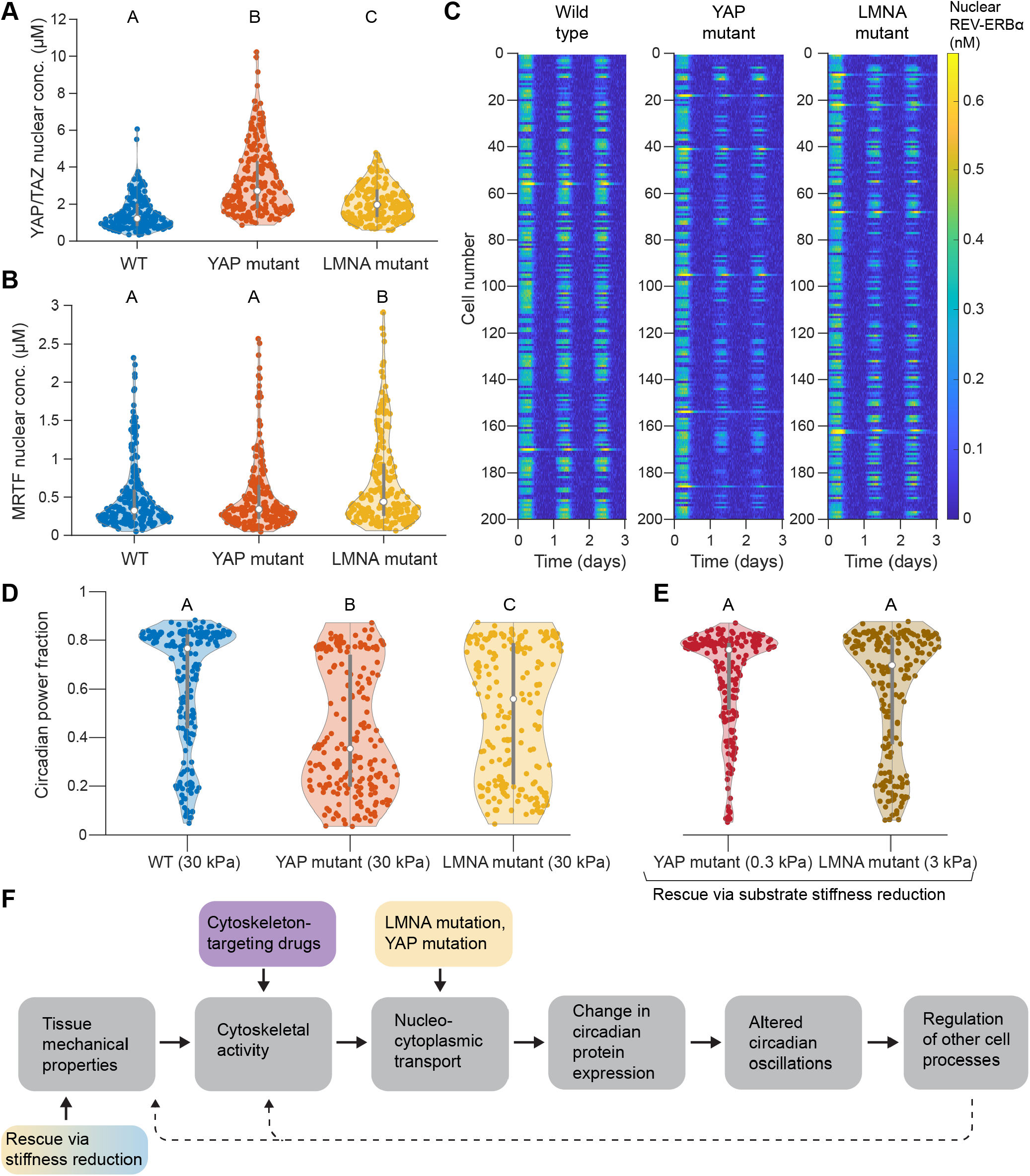
Effects of YAP or lamin A mutations on circadian oscillations. A-B) Nuclear concentrations of YAP/TAZ (A) and MRTF (B) compared across wild type and both mutant populations on 30 kPa substrates. C) Kymograph depicting the dynamics of nuclear REV-ERBα for a wild type population, YAP mutant population, and lamin A mutant population, all on 30 kPa substrates. Each plot shows 200 cells over 5 days. D) Circadian power fraction compared across wild type and mutant cells on 30 kPa substrates. E) Rescue of normal circadian oscillations in mutant cells via reduction of substrate stiffness. Central points and error bars in violin plots denote the median and interquartile range for populations of 200 cells each. Compact letter display is used to denote statistical significance, where groups sharing a letter are statistically similar according to ANOVA followed by Tukey’s *post hoc* test with a significance threshold of *p* = 0.05. Violin plots were generated using Violinplot in MATLAB (Bechtold 2016). F) Summary of the causal flow from cell mechanotransduction to changes in the circadian clock. Upper colored boxes denote disruptions to mechanotransduction explored at different points in this paper, either due to cytoskeleton-targeting drugs or mutations in YAP or LMNA. The lower box depicts the rescue of normal circadian oscillations in mutant cells via a reduction in substrate stiffness. Dashed arrows indicate possible feedback from the circadian clock to cell and tissue mechanics.

Separately, we tested possible effects of mutations in LMNA, the gene controlling the expression of lamin A. We consider a combination of two alterations to the model due to this mutation, aiming to mimic states relevant to laminopathies. Given that laminopathy-associated lamin mutants show distinct differences in their phosphorylation states (Torvaldson et al. 2015; Cenni 2005; Mitsuhashi et al. 2010; Swift et al. 2013), we first assumed that the mutated lamin A was constitutively active (dephosphorylated). We additionally assumed that mutations in lamin A could result in increased nuclear permeability due to changes in localization of nuclear pore complexes (NPCs) or in overall nuclear shape (Dutta et al. 2018). We set the lamin A phosphorylation rate to zero and doubled the opening rate of NPCs (Table S5), leading to increased levels of both YAP/TAZ and MRTF in the nucleus (Figure 6A-B), in agreement with recent experimental data (Owens et al. 2020). Although the effects of such a mutation were informed by the above studies, this condition is not meant to directly represent any one given lamin A mutation, but rather to predict the effects of some possible lamin-related defects. We ensured the robustness of our results by testing different choices of mutation-related parameters for LMNA and 5SA-YAP, as shown in Figure S5.

On 30 kPa substrates, both mutants show altered BMAL1 dynamics, with significantly weaker oscillations compared to wild type cells (Figure 6C). The circadian power fraction decreases distinctly in each case, with the effect being relatively stronger for 5SA-YAP cells (Figure 6D). This is expected, given the dramatic increase in nuclear YAP/TAZ due to overexpression of 5SA-YAP, compared to the milder increases in both YAP/TAZ and MRTF observed for the lamin mutant (Figure 6A-B). This strong influence of 5SA-YAP overexpression on circadian power fraction agrees qualitatively with the findings in Abenza et al. 2023; for ease of comparison, we directly compare our kymographs and circadian power fractions with those from experiments in Figure S6.

Our previous population-level tests establish that lower amounts of mechanical activation (e.g., lower substrate stiffness, treatment with cytoskeletal inhibitors) generally leads to increases in the circadian power fraction (Figure 4, Figure 5). Accordingly, we tested whether a reduction in substrate stiffness might rescue circadian oscillations for either mutant. Remarkably, we find that decreasing substrate stiffness to 3 kPa for the lamin A mutant, or to 0.3 kPa for the 5SA-YAP mutant, results in a power fraction statistically similar to wild type cells on 30 kPa (Figure 6E). This provides a prediction that can readily be tested experimentally. Furthermore, this may indicate a possible compensatory mechanism to counteract perturbations to circadian oscillations in laminopathies.

## Discussion

Circadian rhythms are exhibited by most organisms on earth and appear across biological scales, from oscillations in gene expression in single cells to changes in organism-wide features such as body temperature. These changes in turn regulate physiological processes such as sleep-wake cycles and metabolic activity in healthy organisms. However, disruptions of regular circadian oscillations are associated with various diseases from cancer (Shafi and Knudsen 2019) to diabetes (Mason et al. 2020), placing a high priority on research efforts to understand the regulation of circadian rhythms from the cell to the organism level.

Several recent studies have found that local environmental cues such as substrate stiffness and cytoskeletal activity can modulate the cell circadian clock (Abenza et al. 2023; Xiong et al. 2022; Williams et al. 2018; Yang et al. 2017). However, previous models of circadian oscillations in single cells do not include any effects on the expression of circadian proteins due to such factors. In our minimal model, we propose that nuclear YAP/TAZ and MRTF can alter the expression of BMAL1, PER/CRY, and REV-ERBα independently of other components of the circadian clock. As supported by recent experimental work (Abenza et al. 2023; Xiong et al. 2022), this change in expression seems to be mediated by a combination of TEAD (in the case of YAP/TAZ) and SRF (in the case of MRTF). Although many of the mechanistic details behind this mechanotransduction-circadian coupling remain unclear, our calibrated model fits well to the experimental data and agrees with experimental measurements of cell populations. This suggests that YAP/TAZ and MRTF integrate different mechanical effects on cells, including actin polymerization, cytosolic stiffness, and myosin activity, to provide specific regulatory cues to the circadian clock. These mechanosensitive effects could well extend to other circadian proteins such as RORc and likely differentially regulate different isoforms of PER and CRY. Such details are neglected in the minimal modeling approach we adopt here, but could be well-addressed by integrating future experimental results with more detailed models.

From the main experimental work we draw from here (Abenza et al. 2023; Xiong et al. 2022), the individual roles of YAP/TAZ and MRTF-mediated mechanotransduction remain unclear. While Xiong et al. 2022 focused exclusively on MRTF-mediated activation of SRF, leading to changes in the expression of circadian proteins, Abenza et al. 2023

suggested that YAP/TAZ mediates changes in both BMAL1 and PER/CRY expression. In fact, it was shown in (Abenza et al. 2023) that YAP/TAZ nuclear levels correlate with the circadian power fraction, but MRTF nuclear levels do not. Results from our simulations indicate that these findings need not be contradictory. Although we assume that both YAP/TAZ and MRTF both mediate disruptions to the circadian clock, we find that YAP/TAZ correlates more strongly with the circadian power fraction (Figure 5). However, unlike in Abenza et al. 2023, we do observe a significant correlation between MRTF nuclear to cytosolic ratio and circadian power fraction. This difference appears to be related to the assumed effect of substrate stiffness on the circadian power fraction; our model predicts a larger increase in power fraction for cells on soft substrates than that observed in Abenza et al. 2023. Indeed, if we assume that a separate mechanism leads to increased expression of BMAL1 and PER/CRY in cells on soft substrates, the correlation with MRTF nuclear abundance is no longer significant, whereas the correlation with nuclear YAP/TAZ remains significant (Figure S4).

The overall trend of weaker circadian oscillations for increased mechanical activation agrees well with previous measurements in epithelial cells, but the same study measured the opposite trend in mouse fibroblasts (Williams et al. 2018). This apparently disagrees with the trend observed in mouse fibroblasts by Xiong et al. (Xiong et al. 2022). However, we note that the study by Williams et al. only assessed cell responses to softer hydrogels in 3D culture (Williams et al. 2018), whereas Xiong et al. made all measurements on 2D substrates (Xiong et al. 2022). Aside from tissue- or organism-specific differences in fibroblast behavior, the difference in observations could be attributed to the well-established differences between cell behavior in 2D vs. 3D culture (Baker and Chen 2012; Duval et al. 2017). This could be investigated in future modeling work examining the effects of different spatial stimuli on cell mechanotransduction (Scott et al. 2021).

Our model predicts that certain mutations in YAP/TAZ or lamin A could significantly disrupt circadian oscillations in cell populations (Figure 6). In these cases, increased nuclear accumulation of YAP/TAZ and/or MRTF lead to perturbations to the circadian network such that oscillations are much weaker. This agrees well with experimental measurements in which overexpression or knockout of lamin A leads to significant changes in the expression of some circadian proteins (Roskell 2019). Furthermore, these findings have natural implications for laminopathies, diseases that are characterized by mutations in LMNA. We specifically considered effects of LMNA mutations matching those previously found for laminopathy-associated mutations, including defects in lamin phosphorylation (Torvaldson et al. 2015) and changes in NPC localization and nuclear shape (Dutta et al. 2018). Therefore, disruptions to the cell circadian clock could be one underappreciated component of disease state in laminopathies, as also posited in other recent work (Briand and Collas 2018; Roskell 2019). For instance, due to the well-known role of the circadian clock in regulating metabolic processes (Sahar and Sassone-Corsi 2012), any such disruptions to the circadian clock could contribute to altered metabolism commonly observed in laminopathy patients (Decaudain et al. 2007; Guúenantin et al. 2014).

Our model may provide insights into other disease states as well. For instance, tissue stiffening is a common hallmark of diseases from cancer to atherosclerosis (Chin et al. 2016; Palombo and Kozakova 2016). As predicted by our model (Figure 4), increased substrate stiffness could locally perturb circadian oscillations in cells. Given the host of cellular processes regulated by circadian cues, any such disruptions could have large effects on overall cell function. Additionally, the results of our *in silico* rescue experiment (Figure 6F) suggest that changes in the local mechanical properties of tissue could serve to counteract disruptions to normal circadian oscillations in diseased cells. For instance, a 10-fold reduction (from 30 kPa to 3 kPa) is sufficient to rescue normal circadian oscillations in our simulated LMNA mutant. This may provide an inherent biological strategy to recover normal oscillations in diseases such as laminopathies. Overall, these findings suggest that mutations affecting nuclear transport render cells especially vulnerable to dysfunctions in circadian oscillations upon tissue stiffening.

Other factors currently not considered in our model could have important effects on the nuclear translocation of YAP/TAZ and MRTF, thereby influencing any downstream disruptions to circadian oscillations. For instance, the disruptive effect of substrate stiffness on circadian oscillations could be slightly alleviated for cells in 3D culture conditions, as previous modeling work shows that cells simulated in 2D culture can exhibit higher YAP/TAZ N/C compared to cells in 3D culture (Scott et al. 2021). Moreover, changes in cell shape are known to influence YAP/TAZ nuclear translocation (Scott et al. 2021; Eroumée et al. 2021). Finally, many other pathways such as calcium signaling cascades are known to play important roles in determining the YAP/TAZ N/C (Khalilimeybodi et al. 2023). These various factors provide natural areas for future extensions of our model.

As it stands, the coupling in our model goes only one way; that is, YAP/TAZ and MRTF alter the expression of circadian proteins. In reality, changes to the circadian oscillation period may induce other changes in cell physiology which could couple back to the mechanical state of cells and/or tissue, creating a closed feedback loop (Figure 6F).

For instance, the circadian clock plays an important regulatory role in ECM remodeling, controlling processes from collagen deposition to MMP activity (Dudek et al. 2023; Hahn and Sundar 2023). Rather than assuming the cell reaches mechanical steady state prior to the start of the simulation, YAP/TAZ and MRTF dynamics could readily be integrated within our equations. In the future, this approach would give us a unique opportunity to probe the complex interplay between circadian clocks, mechanotransduction, and disease states.

## Methods

### Numerical implementation

Our code is implemented in MATLAB 2023, and is freely available on Github (**emmeta.francisRangamaniLabUCSDMec** Given the set of treatment conditions (substrate stiffness, cell density, inhibitor concentrations), we algebraically solved for the steady-state concentrations of nuclear YAP/TAZ and nuclear MRTF. These values were then used to compute *K*_*eB*2_, *K*_*eP*2_, and *K*_*eR*2_ (Equations (6) to (8)), and the resulting system of DDEs (Equations (9) to (11)) was solved using dde23. As initial conditions in dde23, we used *B*_0_ = 5*B*^*^, *P*_0_ = 0.2*P* ^*^, and *R*_0_ = *R*^*^, where *B*^*^, *P* ^*^, and *R*^*^ were the steady-state BMAL1, PER/CRY, and REV-ERBα values associated with a given treatment condition. The luciferase concentration, *L*(*t*), was then computed by integrating Equation (12) using ode15s, with the initial condition *L*_0_ = 0.

### Sensitivity analysis and parameter estimation

We conducted a global parametric sensitivity analysis using the UQLab framework in MATLAB (Marelli and Sudret 2014). Treating the luciferase oscillation period and amplitude as quantities of interest, we estimated the total-order Sobol’ indices associated with parameters from Tables S1 and S4. These indices represent the total effect of a parameter on the given output, including first-order effects and all higher-order effects arising from interactions with other parameters. We only considered the case of untreated cells on glass (10 GPa stiffness) and did not test the sensitivity to any inhibitor-related parameters. Sobol’ indices were computed using the default Monte Carlo estimators in UQLab with a sample size of 50,000. Marginal distributions were uniform for all parameters over the fitting ranges specified in Tables S1 and S4.

We completed our model calibration using Bayesian parameter estimation in UQLab, yielding likelihood distributions for each parameter given the mean experimental measurements and associated errors. After computing the luciferase reporter dynamics as described above, we calculate a likelihood function by computing the product of error distributions at discrete time steps for each test. With **p**^*Circ*^ as a vector of free parameters, *k* as the number of experimental conditions, and *N*_*i*_ as the number of time points evaluated for test *i*, the log-likelihood function log(ℒ_*Circ*_) is:

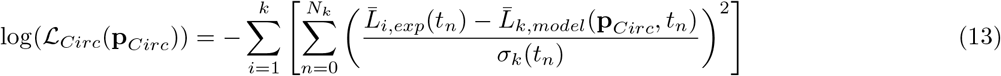

Where 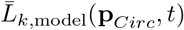 and 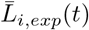 are the normalized luciferase dynamics in the model and experiments. For comparison to the experimental data, we time-shifted *L* to start at the second peak in the simulated dynamics (*t*_*peak*_), normalized to the amplitude of oscillations in the control case (*A*_*model*,*control*_), and subtracted the mean value of *L* for the current condition:

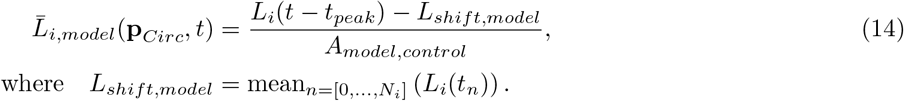

The experimental dynamics were reconstructed from the reported measurements of the period and amplitude of luminescence oscillations in Xiong et al. 2022. For each condition, Xiong et al. measured the population average and standard deviation for the period and amplitude of oscillation (Table S6). Treating the period (*T*_*i*,*exp*_) and amplitude (*A*_*i*,*exp*_) as normally distributed random variables, the relative luminescence over time is given by:

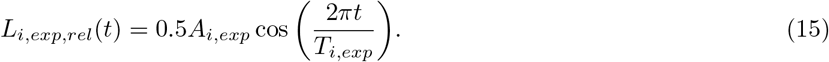

We generate instances o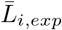 fby sampling from the distributions for *T*_*i*,*exp*_ and *A*_*i*,*exp*_ for each condition. We assume the period and amplitude are normally distributed with mean and standard deviations from Table S6 and generate 1 million pairs of samples per condition. The mean 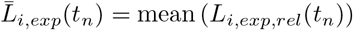 and standard devation *σ*_*i*_(*t*_*n*_) = std (*L*_*i*,*exp*,*rel*_(*t*_*n*_)) for the luminescence were then computed at designated time points.

To better match the experimentally measured oscillation periods, we found it necessary to augment the above log-likelihood function by adding an extra penalty to any model deviations from the experimentally measured oscillation periods:

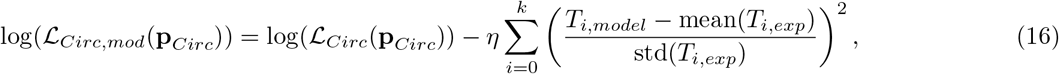

where *η* is a weighting factor determining the magnitude of the penalty term. We tested different values of this factor, finding good performance for *η* = 100.

We sample from the posterior distribution using Markov chain Monte Carlo (MCMC) in UQLab. Specifically, we use the built-in affine invariant ensemble algorithm with 60 walkers and 1,000 total steps. Prior distributions were chosen to be uniform over the ranges specified in Tables S1 and S4. We assessed convergence of MCMC using the integrated autocorrelation time (IACT), which corresponds to the number of steps in the MCMC algorithm required for all walkers to decorrelate from their initial trajectory. We computed this using previously developed code (Wolff 2004; Linden et al. 2022), finding an approximate value of 88 steps. As a conservative measure, we discard the first 500 steps as burn-in when sampling from the posterior distributions. The posterior distributions for each parameter are plotted in Figure S2C.

### Linear stability analysis

To assess which parameters naturally give rise to sustained circadian oscillations, we conducted a linear stability analysis. Numerical bifurcation analysis was conducted using DDE-BIFTOOL in MATLAB (Engelborghs et al. 2002). For this analysis, we used the maximum *a posteriori* (MAP) values as point estimates for each circadian parameter (Table S1) and treated the nuclear to cytosolic ratios of YAP/TAZ and MRTF as bifurcation parameters.

In this context, the Hopf bifurcation is defined by the transition of the rightmost (most positive / least negative) pair of eigenvalues from negative real values to positive real values (Figure S3A). This corresponds to a supercritical Hopf bifurcation, in which damped oscillations transition to a stable limit cycle.

Given the known importance of degradation rates on circadian oscillations, we also examined the effects of *K*_*dB*_, *K*_*dP*_, and *K*_*dR*_ on the location of Hopf bifurcations and the period of oscillations (Figure S3C-E).

### Equations for inhibitor treatments

Throughout this work, we considered several different treatments known to modify the cytoskeleton via distinct mechanisms. Both cytochalasin D and latrunculin A/B have an overall inhibitory effect on actin polymerization, but they trigger distinct effects on MRTF vs. YAP/TAZ nuclear transport (Finch-Edmondson and Sudol 2016; Miralles et al. 2003), prompting careful consideration in our model.

Under standard conditions, actin polymerization can be written in the following form (see Tables S2 and S3 for full details):

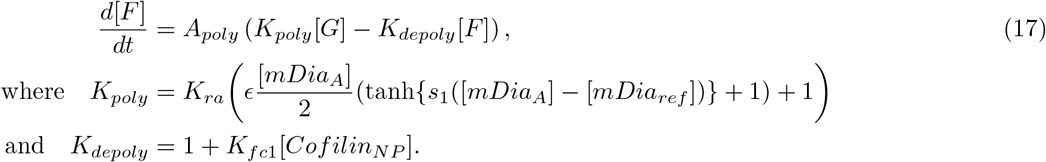

[*F*] is the concentration of F-actin and [*G*] is the concentration of G-actin. *A*_*poly*_ is a constant that sets the overall timescale, which can be arbitrarily set for the sake of the steady-state analysis here. Note that the concentration of [*F*] corresponds to the concentration of actin monomer units per volume; to convert this to the molecular concentration of filaments we would divide by the average length of actin filaments in the cell. Taking into account mass conservation ([*F*] + [*G*] = [*Actin*_*tot*_]), the steady state concentration of [*F*] is:

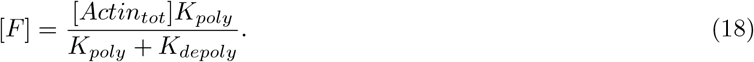

To model the action of cytochalasin D and latrunculin A/B, we use a simplified version of the equations from Wakatsuki et al. 2001. In all cases, we assume that inhibitor is present in excess.

### Cytochalasin D treatment

Cytochalasin D acts by capping existing actin filaments and inducing dimerization of G-actin. Accordingly, we consider additional differential equations for cytochalasin D-induced dimerization ([*CG*_2_]) and cytochalasin D capping of F-actin ([*FC*]):

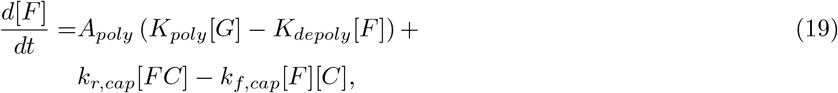

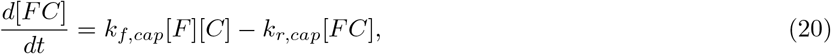

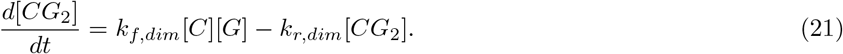

Note that, for simplicity, we assume that the rate of dimerization scales with [*C*][*G*] rather than [*C*][*G*]^2^. This is equivalent to assuming that all cases of cytochalasin D / G-actin binding result in G-actin dimerization after the first binding event, capturing the same qualitative effect while yielding a simpler closed form expression for [*F*]. The corresponding steady-state concentration of F-actin can be readily solved for, assuming mass conservation ([*F*] + [*G*] + 2[*CG*_2_] + [*FC*] = [*Actin*_*tot*_]):

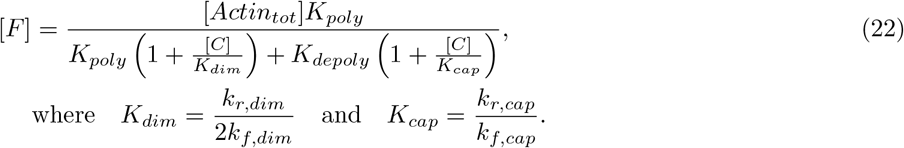

We assume that dimerized G-actin in the case of cytochalasin D treatment no longer sequesters MRTF. Therefore, the relevant G-actin concentration [G] is given by:

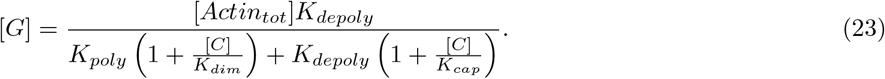

### Latrunculin A/B treatment

Latrunculin A and B act by sequestering G-actin. Writing latrunculin concentration as [*L*] and the G-actin-latrunculin complex concentration as [*GL*], we have the following set of ODEs:

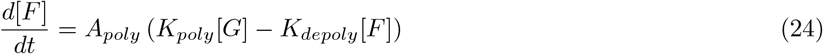

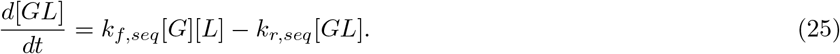

Accounting for conservation of mass ([*G*] + [*GL*] + [*F*] = [*Actin*_*tot*_]), the steady state solution for F-actin and total G-actin can be written as:

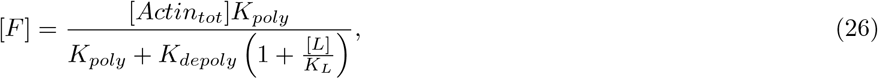

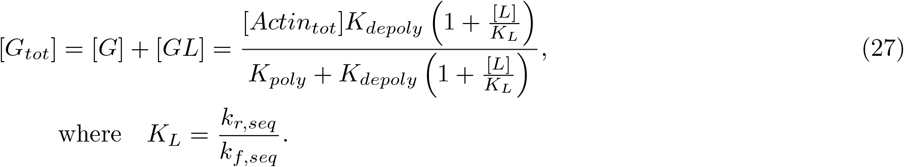

Note that the total G-actin [*G*_*tot*_] is assumed to be the relevant concentration for MRTF sequestration and is therefore the quantity used to compute the amount of free MRTF in this case (see Table S2).

### Jasplakinolide treatment

In the case of Jasplakinolide treatment, in line with in vitro data (Bubb et al. 2000), we made the empirical assumption that actin polymerization rate increases as a function of Jasplakinolide concentration [*J*]:

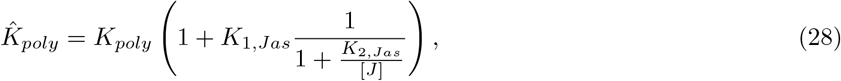

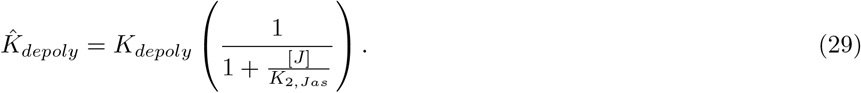

Note that in the general case, 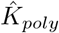 and 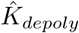 could have different sensitivities to jasplakinolide (*K*_2,*Jas*_ here), but existing data supports these sensitivities being similar (Bubb et al. 2000). In this case, the new steady-state F-actin concentration is:

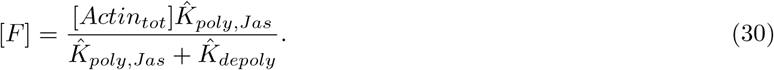

### Other inhibitor treatments and contact area considerations

For all other inhibitor treatments, we assume general Hill equations for concentration-dependent inhibition. The effects of all inhibitor treatments and mutations are summarized in Table S5.

In all cases from Abenza et al., we additionally considered the effects of changes in cell-substrate contact area. We approximated contact area by tracing out the fluorescence microscopy images in Figure 3D and Figure S3A of their paper (Abenza et al. 2023). Control cells had contact area *A*_*contact*,0_ *≈*3000 µm^2^, and any changes in contact area were captured by modulating the amount of FAK phosphorylation:

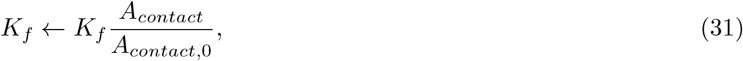

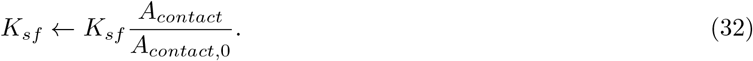

### Simulation of cell populations

Model populations of cells were generated by sampling parameter values from probability distributions. By default, all parameters in our YAP/TAZ mechanotransduction model were sampled from log-normal distributions. We selected parameter values from each distribution by first generating a normally distributed random number *r* using randn() in MATLAB, and then computing:

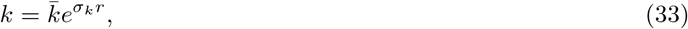

where *σ*_*k*_ is the standard deviation associated with the log-normal distribution of values for *k* and 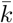 is the baseline value of the parameter. A value of *σ* = 0.2 was used for all parameters except for exponents, which were kept constant.

For all MRTF and circadian parameters fit using the Bayesian approach described above, their probability distributions were described by their posterior distributions (Figure S2C). We randomly drew from these distributions by sampling from the MCMC chains after discarding the first 500 iterations as burn-in.

### Calculation of the circadian power fraction

We used the circadian power fraction as a metric for the quality of circadian oscillations, as defined by Abenza et al. (Abenza et al. 2023). In their paper, they measured circadian oscillations using the fluorescent sensor REV-VNP (Nagoshi et al. 2004) as an indicator for REV-ERBα expression, and so we compute the power fraction from our predictions of REV-ERBα dynamics. Before computing the power fraction, we added white Gaussian noise to individual REV-ERBα trajectories, assuming a signal-to-noise ratio of 5. We sampled the resulting signal every 15 minutes (sampling frequency *f*_*S*_ = 96 day^*−*1^) and then filtered it using a low pass filter with the same settings as in Abenza et al. (filtfilt() in MATLAB with a = 1 and b = (0.2, 0.2, 0.2, 0.2)) and scaled the dynamics by subtracting the mean and dividing by the standard deviation. We then computed the power spectrum, 𝒫, for each cell in the model population for a given condition. The power fraction for an individual cell was then given by:

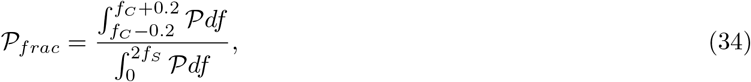

where *f*_*C*_ is the frequency associated with the maximum population-wide average power in the interval 0.7 day^*−*1^ to 1.3 day^*−*1^.

## Acknowledgements

We thank Nathaniel Linden for valuable input on Bayesian parameter estimation, as well as Dr. Ali Khalilimeybodi and Maríia Hernáandez Mesa for their close reading of the manuscript and helpful suggestions.

## Contributions

Conceptualization: E.A.F., P.R.; Methodology: E.A.F., P.R.; Software: E.A.F.; Formal analysis: E.A.F.; Resources: P.R.; Investigation: E.A.F., Data curation: E.A.F.; Writing - original draft: E.A.F.; Writing - review editing: E.A.F. and P.R.; Visualization: E.A.F.; Supervision: P.R.; Project administration: P.R.; Funding acquisition: E.A.F. and P.R.

## Data Availability

Our code is freely available on Github and is deposited on Zenodo (Francis 2024b); all results files are also deposited on Zenodo (Francis 2024a).

