## Supplementary Tables and Figures for "Computational modeling establishes mechanotransduction as a potent modulator of the mammalian circadian clock"

Table S1: Circadian clock model parameters.

| Parameter | Units | Description | Baseline | Source | Fitting range (%) | MAP |
|---|---|---|---|---|---|---|
| $\tau_B$ | hr | BMAL1 delay constant | 12.0 | Tamaru et al. 2003 | 50-200 | 17.75 |
| $K_{eB0}$ | $hr^{-1}$ | BMAL1-dependent BMAL1 expression rate | 0.01 | * | 25-400 | 0.03564 |
| $K_{iR}$ | 1 | Inhibition constant for BMAL1 expression | 0.04 | Lema et al. 2000 | 25-400 | 0.08309 |
| $n_B$ | 1 | Hill coefficient for BMAL1 expression | 2 | Sriram et al. 2006 | 50-200 | 2.223 |
| $K_{dB}$ | $hr^{-1}$ | BMAL1 degradation rate | 0.4 | Lema et al. 2000 | 25-400 | 0.9152 |
| $\tau_P$ | hr | PER/CRY delay constant | 7.5 | Yagita et al. 2001 Yagita et al. 2002 Lee et al. 2001 | 50-200 | 10.59 |
| $K_{eP0}$ | $hr^{-1}$ | BMAL1-dependent PER/CRY expression rate | 0.1 | * | 25-400 | 0.2149 |
| $K_{aB}$ | 1 | Activation constant for PER/CRY expression | 0.5 | * | 25-400 | 0.5068 |
| $n_{P0}$ | 1 | First Hill coefficient for PER/CRY expression | 2 | Lema et al. 2000 | 50-200 | 1.415 |
| $K_{eP1}$ | $hr^{-1}$ | PER/CRY-dependent PER/CRY expression rate | 1.0 | * | 25-400 | 3.835 |
| $K_{iP}$ | 1 | Inhibition constant for PER/CRY expression | 0.1 | * | 25-400 | 0.02981 |
| $n_{P1}$ | 1 | Second Hill coefficient for PER/CRY expression | 2 | Sriram et al. 2006 Lema et al. 2000 | 50-200 | 3.362 |
| $K_{dP}$ | $hr^{-1}$ | PER/CRY degradation rate | 0.4 | * | 25-400 | 0.9478 |
| $\tau_R$ | hr | REV-ERB$\alpha$ delay constant | 7.5 | Sriram et al. 2006 | 50-200 | 9.075 |
| $K_{dR}/\zeta$ | $hr^{-1}$ | REV-ERB$\alpha$ degradation rate (relative) | 0.4 | * | 25-400 | 0.6888 |
| $K_{eB2,Y}$ | $hr^{-1}$ | YAP/TAZ-regulated BMAL1 expression rate | 0.05 | * | 25-400 | 0.1328 |
| $K_{YB}$ | nM | Activation constant for YAP/TAZ regulation of BMAL1 expression | 1.0 | * | 25-400 | 3.109 |
| $K_{eB2,M}$ | $hr^{-1}$ | MRTF-regulated BMAL1 expression rate | 0.05 | * | 25-400 | 0.1609 |
| $K_{MB}$ | μM | Activation constant for MRTF regulation of BMAL1 expression | 1.0 | * | 25-400 | 2.298 |
| $K_{eP2,Y}$ | $hr^{-1}$ | YAP/TAZ-regulated PER/CRY expression rate | 0.05 | * | 25-400 | 0.03885 |
| $K_{YP}$ | μM | Activation constant for YAP/TAZ regulation of PER/CRY expression | 1.0 | * | 25-400 | 2.001 |
| $K_{eP2,M}$ | $hr^{-1}$ | MRTF-regulated PER/CRY expression rate | 0.05 | * | 25-400 | 0.1693 |
| $K_{MP}$ | μM | Activation constant for MRTF regulation of PER/CRY expression | 1.0 | * | 25-400 | 2.309 |
| $K_{eR2,Y}$ | $hr^{-1}$ | YAP/TAZ-regulated REV-ERB$\alpha$ expression rate | 0.05 | * | 25-400 | 0.1356 |
| $K_{YR}$ | μM | Activation constant for | 1.0 | * | 25-400 | 1.273 |

Continued on next page

**Table S1:** (continued.) Circadian clock model parameters.

| Parameter | Units | Description | Baseline | Source | Fitting range (%) | MAP |
|---|---|---|---|---|---|---|
| | | YAP/TAZ regulation of REV-ERB$\alpha$ expression | | | | |
| $K_{eR2,M}$ | hr$^{-1}$ | MRTF-regulated REV-ERB$\alpha$ expression rate | 0.05 | * | 25-400 | 0.1024 |
| $K_{MR}$ | µM | Activation constant for MRTF regulation of REV-ERB$\alpha$ expression | 1.0 | * | 25-400 | 1.737 |
| $K_{dL}$ | hr$^{-1}$ | Luciferase degradation rate | 0.35 | Feeney et al. 2016 | 25-400 | 0.1341 |
| $\tau_L$ | hr | Luciferase delay constant | 1 | ** | ** | 1 |
| $B_{ref}$ | nM | Reference BMAL1 concentration | 190 | † | † | 190 |
| $P_{ref}$ | nM | Reference PER/CRY concentration | 7.5 | † | † | 7.5 |
| $R_{ref}$ | nM | Reference REV-ERB$\alpha$ concentration | 0.07 | † | † | 0.07 |

Fitting range specifies the range of the uniform prior distribution (in percentage) and "MAP" is the maximum *a posteriori* from Bayesian parameter estimation.

* Set based on early testing of the model, such that oscillations still occur when carrying over applicable parameter values from Lema et al. 2000.

** The choice of $\tau_L$ is arbitrary in this equation; it does not affect model behavior, as it simply sets the phase of luciferase oscillations, but changes in luceriferase do not feedback to affect any other dynamics.

† To convert dimensionless BMAL1, PER/CRY, and REV-ERB$\alpha$ concentrations to nM, we multiplied these concentrations by $B_{ref}$, $P_{ref}$, and $R_{ref}$, respectively. We set these parameters to match with experimental measurements (Smyllie et al. 2016) and previous models (Mirsky et al. 2009). These parameters are fixed and not used in fitting as our fitting protocol only examines relative changes in nuclear concentration (see Supplementary Information).

**Table S2:** Mechanotransduction model equations. The ODE expression and steady-state expression are given for each variable considered. Note that only the steady-state solutions were considered in this study.

| Variable | Expression |
|---|---|
| $\frac{d[FAK_P]}{dt}$ | $\left(k_f + k_{sf}\frac{E}{C_{stiffness}+E}\right)([FAK_{tot}]-[FAK_P]) - k_{df}[FAK_P]$ |
| $[FAK_P]_{SS}$ | $[FAK_{tot}]\frac{K_f(C_{stiffness}+E)+K_{sf}E}{(1+K_f)(C_{stiffness}+E)+K_{sf}E}$ where $K_f = \frac{k_f}{k_{df}}$, $K_{sf} = \frac{k_{sf}}{k_{df}}$ |
| $\frac{d[RhoA_{GTP}]}{dt}$ | $k_{fk\rho}(\gamma[pFAK]^n+1)\left(\frac{[RhoA_{tot}]}{SA_{PM}} - [RhoA_{GTP}]\right) - k_{d\rho}[RhoA_{GTP}]$ |
| $[RhoA_{GTP}]$ | $\frac{[RhoA_{tot}]}{SA_{PM}}\frac{K_{fk\rho}(\gamma[pFAK]^n_{SS}+1)}{1+K_{fk\rho}(\gamma[pFAK]^n_{SS}+1)}$ where $K_{fk\rho} = \frac{k_{fk\rho}}{k_{d\rho}}$ |
| $\frac{d[ROCK_A]}{dt}$ | $k_{r\rho}[RhoA_{GTP}]\frac{([ROCK_{tot}]-[ROCK_A])SA_{PM}}{vol_{cyto}N_{convert}} - k_d[ROCK_A]$ |
| $[ROCK_A]$ | $[ROCK_{tot}]\frac{K_{r\rho}[RhoA_{GTP}]_{SS}SA_{PM}}{vol_{cyto}N_{convert}+K_{r\rho}[RhoA_{GTP}]_{SS}SA_{PM}}$, where $K_{r\rho} = \frac{k_{r\rho}}{k_d}$ |

Continued on next page

**Table S2:** (continued.) Mechanotransduction model equations.

| Variable | Expressions |
|---|---|
| $\frac{d[mDia_A]}{dt}$ | $k_{m\rho}[RhoA_{GTP}]\frac{([mDia_{tot}]-[mDia_A])SA_{PM}}{vol_{cyto}N_{convert}} - k_{dmDia}[mDia_A]$ |
| $[mDia_A]_{SS}$ | $[mDia_{tot}]\frac{K_{m\rho}[RhoA_{GTP}]_{SS}SA_{PM}}{vol_{cyto}N_{convert}+K_{m\rho}[RhoA_{GTP}]_{SS}SA_{PM}}$ <br> where $K_{m\rho}=\frac{k_{m\rho}}{k_{dmDia}}$ |
| $\frac{d[Myo_A]}{dt}$ | $\frac{1}{2}k_{mr}\big(\epsilon[ROCK_A](\tanh\{s_1([ROCK_A]-[ROCK_{ref}])\}+1)+1\big)([Myo_{tot}]-[Myo_A])$ <br> $-k_{dmy}[Myo_A]$ |
| $[Myo_A]_{SS}$ | $[Myo_{tot}]\frac{K_{mr}\big(\epsilon[ROCK_A]_{SS}(\tanh\{s_1([ROCK_A]_{SS}-[ROCK_{ref}])\}+1)+1\big)}{2+K_{mr}\big(\epsilon[ROCK_A]_{SS}(\tanh\{s_1([ROCK_A]_{SS}-[ROCK_{ref}])\}+1)+1\big)}$ <br> where $K_{mr}=\frac{k_{mr}}{k_{dmy}}$ |
| $\frac{d[LIMK_A]}{dt}$ | $\frac{1}{2}k_{lr}\big(\tau[ROCK_A](\tanh\{s_1([ROCK_A]-[ROCK_{ref}])\}+1)+1\big)([LIMK_{tot}]-[LIMK_A])$ <br> $-k_{dl}[LIMK_A]$ |
| $[LIMK_A]_{SS}$ | $[LIMK_{tot}]\frac{K_{lr}\big(\tau[ROCK_A]_{SS}(\tanh\{s_1([ROCK_A]_{SS}-[ROCK_{ref}])\}+1)+1\big)}{2+K_{lr}\big(\tau[ROCK_A]_{SS}(\tanh\{s_1([ROCK_A]_{SS}-[ROCK_{ref}])\}+1)+1\big)}$ <br> where $K_{lr}=\frac{k_{lr}}{k_{dl}}$ |
| $\frac{d[Cofilin_{NP}]}{dt}$ | $k_{turnover}([Cofilin_{tot}]-[Cofilin_{NP}])-\frac{k_{cat,Cof}[LIMK_A][Cofilin_{NP}]}{k_{m,Cof}+[Cofilin_{tot}]}$ |
| $[Cofilin_{NP}]_{SS}$ | root$\left\{k_{turnover}([Cofilin_{tot}]-[Cofilin_{NP}]_{SS})-\frac{k_{cat,Cof}[LIMK_A]_{SS}[Cofilin_{NP}]_{SS}}{k_{m,Cof}+[Cofilin_{tot}]}\right\}$ |
| $\frac{d[F]}{dt}$ | $\frac{1}{2}k_{ra}\big(\alpha[mDia_A](\tanh\{s_1([mDia_A]-[mDia_{ref}])\}+1)+1\big)([Actin_{tot}]-[F])$ <br> $-(k_{dep}+k_{fc1}[Cofilin_{NP}])[F]$ |
| $[F]_{SS}$ | $\frac{[Actin_{tot}]K_{ra}\big(\alpha[mDia_A]_{SS}(\tanh\{s_1([mDia_A]_{SS}-[mDia_{ref}])\}+1)+1\big)}{2(1+K_{fc1}[Cofilin_{NP}]_{SS})+K_{ra}\big(\tau[mDia_A]_{SS}(\tanh\{s_1([mDia_A]_{SS}-[mDia_{ref}])\}+1)+1\big)}$ <br> where $K_{ra}=\frac{k_{ra}}{k_{dep}},\quad K_{fc1}=\frac{k_{fc1}}{k_{dep}}$ |
| $\frac{d[LaminA_{NP}]}{dt}$ | $\frac{k_{fl}E_{cyto}}{C_{LaminA}+E_{cyto}}([LaminA_{tot}]-[LaminA_{NP}])-k_{rl}[LaminA_{NP}]$ |
| $[LaminA_{NP}]_{SS}$ | $[LaminA_{tot}]\frac{K_{fl}E_{cyto}}{C_{LaminA}+E_{cyto}(1+K_{fl})}$, where $E_{cyto}=p_{cyto}[F]^{2.6}$ and $K_{fl}=\frac{k_{fl}}{k_{rl}}$ |

Continued on next page

**Table S2:** (continued.) Mechanotransduction model equations.

| Variable | Expressions |
|---|---|
| $\frac{d[NPC_A]}{dt}$ | $k_{fNPC}[LaminA_{NP}][F][MyoA]([NPC_{tot}]-[NPC_A])-k_r[NPC_A]$ |
| $[NPC_A]_{SS}$ | $[NPC_{tot}]\frac{K_{fNPC}[LaminA_{NP}]_{SS}[F]_{SS}[MyoA]_{SS}}{1+K_{fNPC}[LaminA_{NP}]_{SS}[F]_{SS}[MyoA]_{SS}}$ where $K_{fNPC}=\frac{k_{fNPC}}{k_r}$ |
| $\frac{d[Y_{NP}]}{dt}$ | $(k_{CN}+k_{CY}[F][Myo_A])([Y_{cyto}]-[Y_{NP}])-k_{NC}[Y_{NP}]$ |
| $[Y_{NP}]_{SS}$ | $\frac{[Y_{tot}]-[Y_{nuc}]_{SS}vol_{nuc}N_{convert}}{vol_{cyto}N_{convert}}\phi_{NP}$ where $K_{CN}=\frac{k_{CN}}{k_{NC}}$, $K_{CY}=\frac{k_{CY}}{k_{NC}}$, $\phi_{NP}=\frac{K_{CN}+K_{CY}[F]_{SS}[Myo_A]_{SS}}{K_{CN}+K_{CY}[F]_{SS}[Myo_A]_{SS}+1}$ * |
| $\frac{d[Y_{nuc}]}{dt}$ | $\frac{SA_{NM}}{vol_{nuc}}((k_{inb}+k_{in}[NPC_A])[Y_{NP}]-k_{out}[Y_{nuc}])$ |
| $[Y_{nuc}]_{SS}$ | $\frac{[Y_{tot}]\phi_{NP}}{N_{convert}}\frac{K_{in,solo,Y}+K_{in,2,Y}[NPC_A]_{SS}}{(K_{in,solo,Y}+K_{in,2,Y}[NPC_A]_{SS})\phi_{NP}vol_{nuc}+vol_{cyto}}$ where $K_{inb}=\frac{k_{inb}}{k_{out}}$, $K_{in}=\frac{k_{in}}{k_{out}}$ |
| $\frac{d[M_{nuc}]}{dt}$ | $\frac{SA_{nuc}}{vol_{nuc}N_{convert}}\left((k_{in,solo,MRTF}+k_{in,2,MRTF}[NPC_A])[M_{free}]-k_{out,MRTF}[M_{nuc}]\right)$ where $[M_{free}]=[M_{cyto}]\frac{1}{1+\left(\frac{[G]}{K_{MRTF}}\right)^2}$ and $[G]=[Actin_{tot}]-[F]$ |
| $[M_{nuc}]_{SS}$ | $\frac{[M_{tot}]}{N_{convert}}\frac{K_{in,solo,MRTF}+K_{in,2,MRTF}[NPC_A]_{SS}}{(K_{in,solo,MRTF}+K_{in,2,MRTF}[NPC_A]_{SS})vol_{nuc}+\left(1+\left(\frac{[G]_{SS}}{K_{MRTF}}\right)^2\right)vol_{cyto}}$ where $K_{in,solo,MRTF}=\frac{k_{in,solo,MRTF}}{k_{out,MRTF}}$, $K_{in,2,MRTF}=\frac{k_{in,2,MRTF}}{k_{out,MRTF}}$, and $[G]_{SS}=[Actin_{tot}]-[F]_{SS}$ |

We include the rate of change and the associated steady-state solution for each variable in Scott et al. 2021. Note that we only use the steady-state relationships in our implementation here, which effectively eliminates one parameter in each equation. An "A" subscript denotes an activated form of a species, "P" denotes a phosphorylated form, and "NP" denotes a nonphosphorylated form.
* $\phi_{NP}$ corresponds to the fraction of cytosolic YAP/TAZ that is not phosphorylated.

**Table S3: YAP/TAZ mechanotransduction parameters**

| Parameter | Units | Description | Value |
|---|---|---|---|
| $E$ | kPa | Substrate stiffness | (varies) |
| $K_f$ | 1 | FAK basal activation constant | 0.4286 |
| $K_{sf}$ | 1 | substrate-stiffness-dependent FAK activation constant | 10.83 |
| $K_{fk\rho}$ | 1 | RhoA basal activation constant | 0.02688 |
| $\gamma$ | $\mu M^5$ | pFAK-dependent RhoA activation constant | 77.56 |
| $n$ | 1 | Cooperativity factor between FAK and Rho | 5 |
| $K_{r\rho}$ | $\mu M^{-1}$ | RhoA-dependent ROCK activation constant | 0.8100 |
| $K_{m\rho}$ | $\mu M^{-1}$ | RhoA-dependent mDia activation constant | 0.4000 |
| $K_{mr}$ | 1 | Myosin activation constant | 0.4478 |
| $\epsilon$ | $\mu M^{-1}$ | ROCK-dependent amplification of myosin activation | 36 |
| $s_1$ | $\mu M^{-1}$ | Slope of $\tanh$ activation functions | 20 |
| $[ROCK_{ref}]$ | µM | $[ROCK_A]$ threshold in activation functions | 0.3 |
| $K_{lr}$ | 1 | LIMK activation constant | 0.035 |
| $\tau$ | $\mu M^{-1}$ | ROCK-dependent amplification of LIMK activation | 55.49 |
| $k_{turnover}$ | $s^{-1}$ | Cofilin activation rate | 0.04 |
| $k_{cat,Cof}$ | $s^{-1}$ | Rate of catalysis for cofilin phosphorylation | 0.34 |
| $k_{m,Cof}$ | µM | $[Cofilin_{NP}]$ at half max phosphorylation rate | 4 |
| $K_{ra}$ | 1 | Actin polymerization constant | 0.1143 |
| $\alpha$ | $\mu M^{-1}$ | mDia-dependent amplification of actin polymerization | 50 |
| $[mDia_{ref}]$ | µM | $[mDia_A]$ threshold in activation functions | 0.165 |
| $K_{fc1}$ | $\mu M^{-1}$ | Cofilin-dependent deactivation of actin polymerization | 1.1429 |
| $K_{fl}$ | 1 | Cytosolic stiffness-dependent lamin A activation constant | 460 |
| $C_{laminA}$ | kPa | Threshold stiffness for lamin A activation | 100 |
| $p_{cyto}$ | kPa-$\mu M^{-2.6}$ | Proportionality between cytosolic stiffness and $[F]^{2.6}$ | $9 \times 10^{-6}$ |
| $K_{fNPC}$ | $\mu m^2 \mu M^{-1}$ | NPC opening constant | $3.218 \times 10^{-8}$ |
| $K_{CN}$ | 1 | Baseline YAP/TAZ activation constant | 4.0 |
| $K_{CY}$ | $\mu M^{-2}$ | Stress-fiber-dependent YAP/TAZ activation constant | $5.429 \times 10^{-3}$ |
| $K_{in,solo,Y}$ | 1 | Basal YAP/TAZ nuclear import constant | 1 |
| $K_{in,2,Y}$ | $\mu m^2$ | $[NPC_A]$-dependent YAP/TAZ nuclear import constant | 10 |
| $vol_{cyto}$ | $\mu m^3$ | Cytosolic volume | 2300 |
| $SA_{PM}$ | $\mu m^2$ | Plasma membrane surface area | 1260 |
| $vol_{nuc}$ | $\mu m^3$ | Nuclear volume | 390 |
| $[FAK_{tot}]$ | µM | Total cytosolic $[FAK]$ | 1 |
| $[RhoA_{tot}]$ | molecules | Total RhoA molecules per cell | $1.385 \times 10^6$ |
| $[ROCK_{tot}]$ | µM | Total cytosolic $[ROCK]$ | 1 |
| $[mDia_{tot}]$ | µM | Total cytosolic $[mDia]$ | 0.8 |
| $[Myo_{tot}]$ | µM | Total cytosolic $[Myo]$ | 5 |
| $[LIMK_{tot}]$ | µM | Total cytosolic $[LIMK]$ | 2 |
| $[Cofilin_{tot}]$ | µM | Total cytosolic $[Cofilin]$ | 2 |
| $[Actin_{tot}]$ | µM | Total cytosolic actin (G-actin monomers per volume) | 500 |
| $[LaminA_{tot}]$ | $\#\mu m^{-2}$ | Total lamin A molecules in the nuclear membrane | 3500 |
| $[NPC_{tot}]$ | $\#\mu m^{-2}$ | Total NPCs in the nuclear membrane | 6.5 |
| $[YAPTAZ_{tot}]$ | molecules | Total YAP/TAZ molecules per cell | $1.385 \times 10^6$ |
| $[MRTF_{tot}]$ | molecules | Total MRTF molecules per cell | $1 \times 10^6$ |
| $N_{convert}$ | $(\#/\mu m^3)/\mu M$ | Unit conversion factor | 602.2 |

Table S4: **Estimated MRTF mechanotransduction and inhibitor treatment parameters.**

| Parameter | Units | Description | Baseline | Source | Fitting range (%) | MAP |
|---|---|---|---|---|---|---|
| $K_{in,solo,MRTF}$ | 1 | MRTF nuclear entry rate constant | 1 | Scott et al. 2021 | 25-400 | 1.132 |
| $K_{in,2,MRTF}$ | μm$^2$ | Lamin-A-dependent MRTF nuclear entry rate constant | 10 | Scott et al. 2021 | 25-400 | 7.756 |
| $K_{release,MRTF}$ | μM | Binding affinity of MRTF to G-actin | 100 | * | 25-400 | 219.8 |
| $C_{stiffness}$ | kPa | Threshold stiffness for FAK phosphorylation | 3.25 | Scott et al. 2021 | 25-400 | 4.450 |
| $K_{cap}$ | μM | Cytochalasin D binding affinity for F-actin capping | 2 | † | 25-400 | 5.040 |
| $K_{dim}$ | μM | Cytochalasin D binding affinity for G-actin dimerization | 2 | † | 25-400 | 1.756 |
| $K_{Y27632}$ | μM | Y27632 treatment EC50 | 10 | † | 25-400 | 22.12 |
| $K_L$ | μM | Latrunculin B treatment EC50 | 0.2 | † | 25-400 | 0.3947 |
| $K_{1,Jas}$ | 1 | Jasplakinolide treatment constant 1 | 2 | Bubb et al. 2000 ‡ | 25-400 | 1.123 |
| $K_{2,Jas}$ | μM | Jasplakinolide treatment constant 2 | 0.1 | Bubb et al. 2000 ‡ | 25-400 | 0.09520 |

Fitting range specifies the range of the uniform prior distribution (in percentage) and "MAP" is the maximum *a posteriori* from Bayesian parameter estimation.

* $K_{release,MRTF}$ initialized to a moderate value of G-actin based on the total amount of actin in the cell.

† Moderate treatment concentrations used in Xiong et al. 2022.

‡ Baseline values approximated from observed effects *in vitro*; see Fig 4 in Bubb et al. 2000.

Table S5: **Summary of different treatment effects.**

| **Test** | **Treatment effect on parameters** | $A_{contact}$ * |
|---|---|---|
| Cytochalasin D | $[F] \leftarrow$ Equation 22, $[G] \leftarrow$ Equation 23 | 5000 |
| Latrunculin B | $K_{poly} \leftarrow \frac{K_{poly}}{1+{[LatB]}/{K_L}}$ | 3000 |
| Jasplakinolide | $K_{poly} \leftarrow K_{poly}\left(1+(1+K_{1,Jas})\frac{[J]}{K_{2,Jas}}\right)$ | 3000 |
| Y27632 | $K_{r\rho} \leftarrow \frac{K_{r\rho}}{1+{[Y27632]}/{K_{Y27632}}}$ | 3000 |
| High cell density | $K_{CN} \leftarrow \frac{K_{CN}}{K_{LATS}}, \quad K_{CY} \leftarrow \frac{K_{CY}}{K_{LATS}}$ ** | 1200 |
| Low cell density | (Abenza control) | 3000 |
| 1600 μm$^2$ micropatterns | - | 1600 |
| 900 μm$^2$ micropatterns | - | 900 |
| Latrunculin A | $K_{poly} \leftarrow \frac{K_{poly}}{1+{[LatA]}/{K_L}}$ *** | 600 |
| Blebbistatin | $K_{CY} \leftarrow \frac{K_{CY}}{1+{[bleb]}/{K_{bleb}}}, \quad K_{fNPC} \leftarrow \frac{K_{fNPC}}{1+{[bleb]}/{K_{bleb}}}$ † | 4000 |
| Soft substrate (Abenza) | $E = 0.3$kPa | 1000 |
| 5SA-YAP mutant | $Y_{NP} \leftarrow Y_{NP} + Y_{NP}^{5SA}, Y_{nuc} \leftarrow Y_{nuc} + Y_{nuc}^{5SA}$ where $Y_{NP}^{5SA}$ and $Y_{nuc}^{5SA}$ are $Y_{NP}$ and $Y_{nuc}$ eval. for $K_{CN} \leftarrow \infty$, $K_{CY} \leftarrow \infty$, and $[YAPTAZ_{tot}] \leftarrow K_{overexpress}[YAPTAZ_{tot}]$‡ | § |
| Lamin A mutant | $K_{fl} \leftarrow \infty$ $K_{fNPC} \leftarrow 2K_{fNPC}$ | § |

For population-level models, $K_{dim}$, $K_{cap}$, $K_L$, $K_{1,Jas}$, $K_{2,Jas}$, and $K_{Y27632}$ were all sampled from their posterior distributions. $K_{LATS}$, $K_{bleb}$, $K_{overexpress}$, and $A_{contact}$ were sampled from log-normal distributions centered at their specified baseline values.

* Incorporated through Equations 31 and 32.

** $K_{LATS}$ captures the increased phosphorylation of YAP/TAZ due to LATS activation; its baseline value is 3, to match the maximum LATS-mediated fold increase in YAP/TAZ phosphorylation rate in Sun et al. 2016

*** For simplicity, Latrunculin A is assumed to have the same treatment response as Latrunculin B.

† The baseline value for $K_{bleb}$ is 1 μM.

‡ The baseline value for $K_{overexpress}$ is 1 (same expression level as native YAP/TAZ).

§ Mutant cells are assumed to adopt the same contact areas as wild type cells under a given condition.

Table S6: **Summary of calibration data.**

| Condition | Mean period | Std period | Mean ampl | Std ampl |
|---|---|---|---|---|
| Control (Tot) * | 24.25 | 0.4239 | 1 | 0.3996 |
| Control (glass) | 23.84 | 0.1748 | 1 | 0.3739 |
| 300 kPa substrate | 24.59 | 0.2330 | 2.295 | 0.1094 |
| 19 kPa substrate | 25.59 | 0.3350 | 1.939 | 0.2553 |
| Control (Y27632) | 24.71 | 0.1659 | 1 | 0.0090 |
| 10 μM Y27632 | 25.10 | 0.2341 | 1.306 | 0.1081 |
| 20 μM Y27632 | 25.17 | 0.3024 | 1.153 | 0.1081 |
| Control (CytD) | 24.21 | 0.2060 | 1 | 0.1954 |
| 1 μM CytD | 25.00 | 0.1803 | 1.287 | 0.1494 |
| 2 μM CytD | 24.97 | 0.0258 | 1.655 | 0.3218 |
| 5 μM CytD | 24.64 | 0.3991 | 1.943 | 0.3908 |
| Control (LatB) | 24.94 | 0.1048 | 1 | 0.0235 |
| 1 μM LatB | 25.41 | 0.1921 | 1.612 | 0.1412 |
| 2 μM LatB | 25.97 | 0.2620 | 2.377 | 0.0588 |
| Control (Jas) | 23.68 | 0.2487 | 1 | 0.1250 |
| 0.1 μM Jas | 23.22 | 0.2902 | 0.953 | 0.1797 |
| 0.2 μM Jas | 22.39 | 0.4145 | 1.094 | 0.2344 |
| 0.5 μM Jas | 20.40 | 0.1658 | 0.2969 | 0.0703 |

* Weighted averages of control data (weighted by number of data points associated with each measurement). Standard deviation was computed as the root mean weighted sum of squares (the square root of the mean variance).

### Supplementary Figures

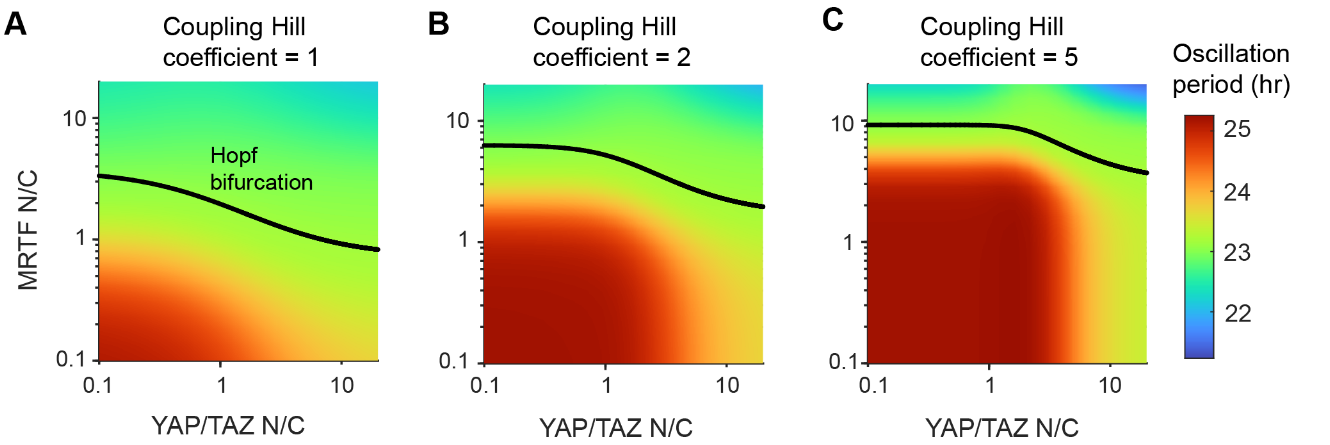

**Figure S1: Effect of Hill coefficient on mechanotransduction-circadian coupling.** Oscillation period and Hopf bifurcation plotted over the MRTF-YAP/TAZ phase plane for different choices of Hill coefficient in Equations 6-8. For simplicity, the same Hill coefficient was used in all equations for each case, either 1 (A), 2 (B, same as all other tests in the paper), or 5 (C). Changing the Hill coefficient simply shifts the Hopf bifurcation line and oscillation period without changing the qualitative behavior of the system.

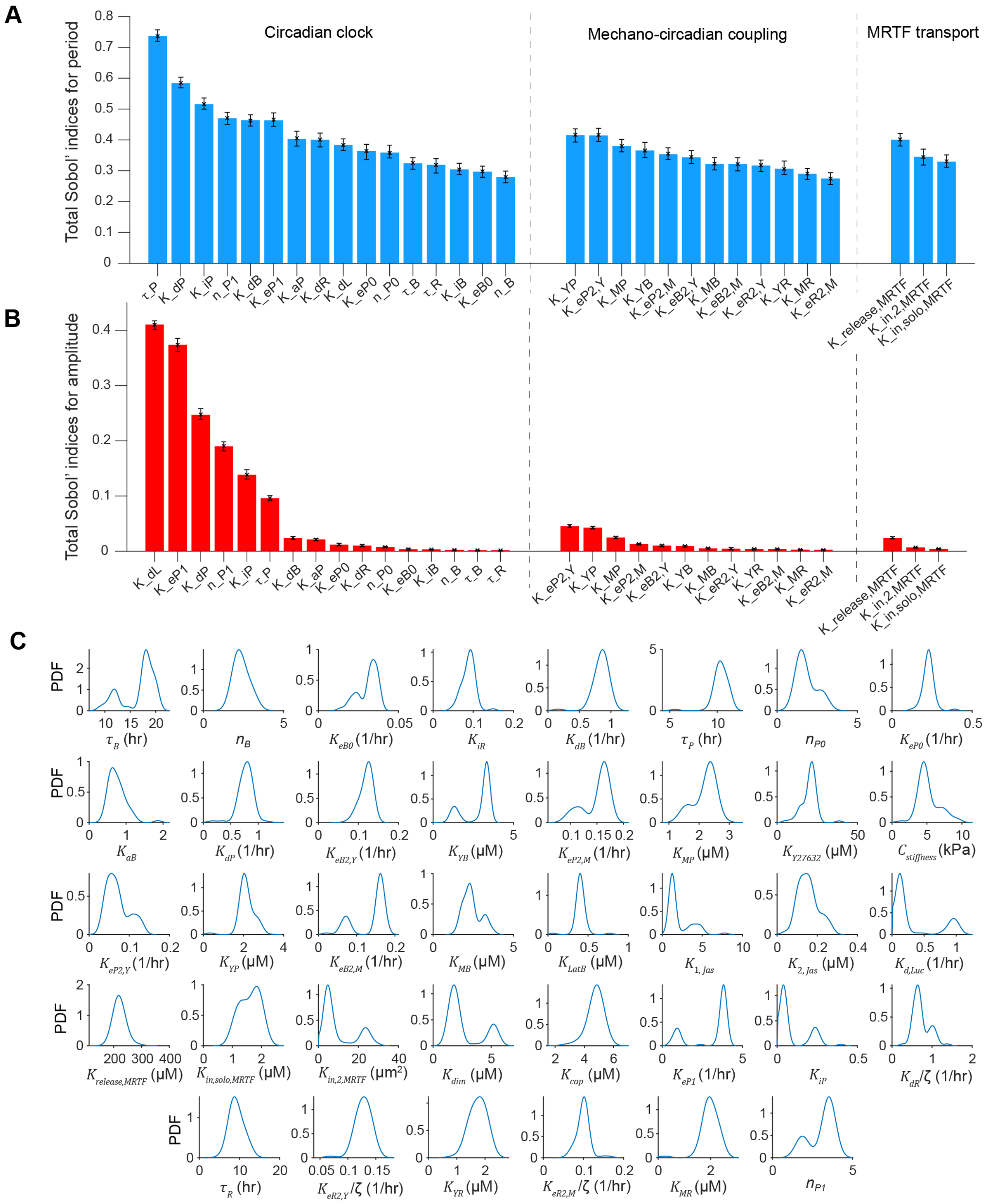

**Figure S2: Sobol' sensitivity analysis and Bayesian parameter estimation.** A-B) Total Sobol' indices associated with the luciferase oscillation period (A) and amplitude (B). Parameters are sorted into three categories: those related exclusively to the core circadian clock, those related to mechanotransduction-circadian coupling, and those governing MRTF transport. Within each category, parameters are plotted from the highest to the lowest estimated total Sobol' index. Markers with error bars denote the mean and 95% confidence intervals computed from bootstrapping with 100 replications. C) Parameter posterior distributions from Bayesian parameter estimation. Markov chain Monte Carlo (MCMC) was run for 1,000 iterations with 60 walkers. After discarding the first 500 iterations as burn-in, posterior distributions were estimated using kernel density estimation in MATLAB, with a bandwidth of 5% of the maximum value of a given parameter included in the prior distribution.

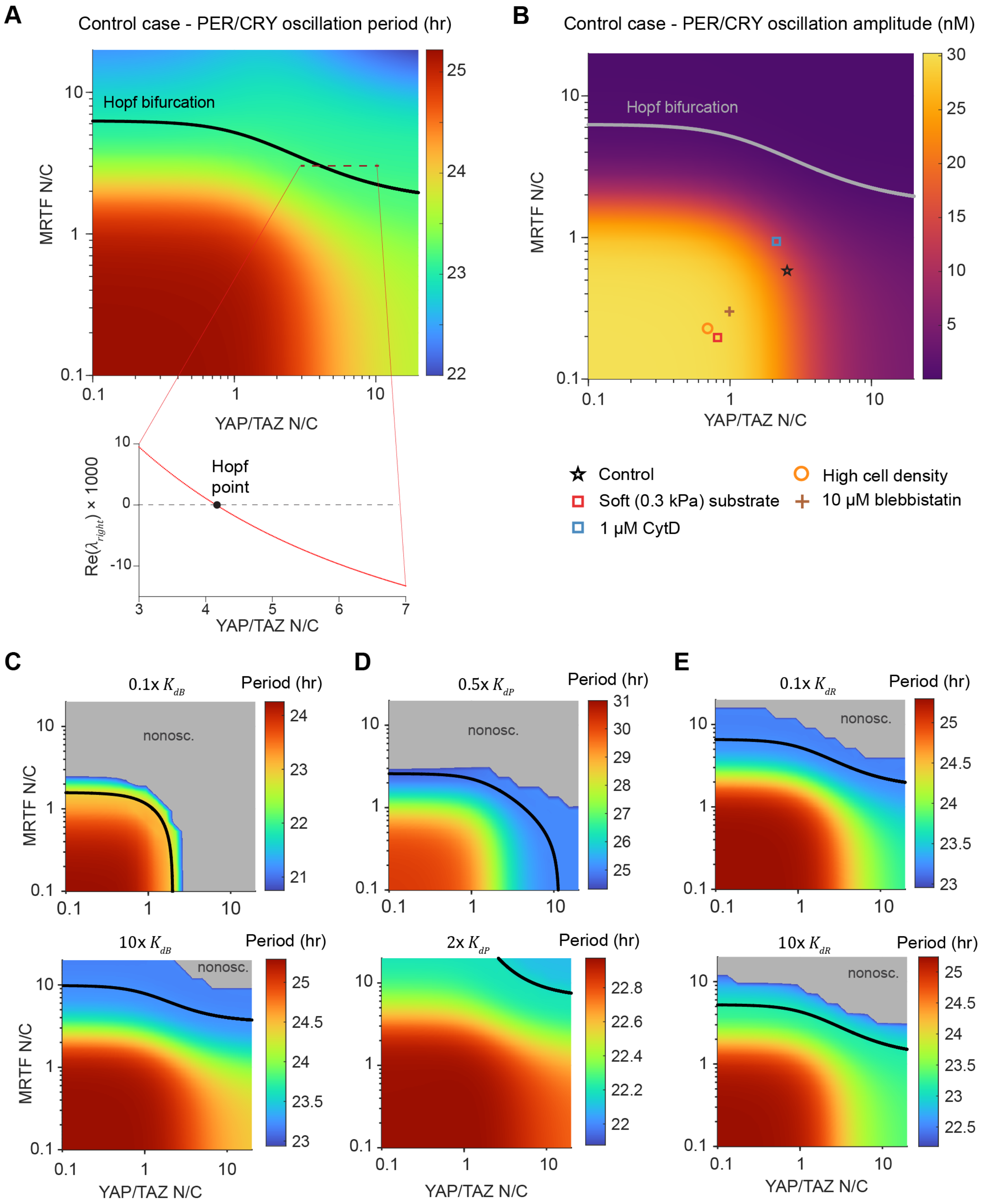

**Figure S3: Linear stability analysis for different parameter values.** A) Bifurcation diagram for control conditions, with oscillation period and location of the Hopf bifurcation plotted as a function of YAP/TAZ and MRTF nuclear to cytosolic ratios (N/C). The real part of the right-most pair of eigenvalues ($\lambda_{right}$) along the line where MRTF N/C is 3 (red dashed line in upper plot). The transition from positive to negative real part of the eigenvalues marks the occurrence of a Hopf bifurcation. B) Oscillation amplitude of PER/CRY over the MRTF-YAP/TAZ phase plane. Individual points correspond to tests run in Figure 5. C-E) Effects of decreasing or increasing $K_{dB}$ (C), $K_{dP}$ (D), or $K_{dR}$ (E) on the location of the Hopf bifurcation and the oscillation period. The grey regions denote nonoscillatory solutions.

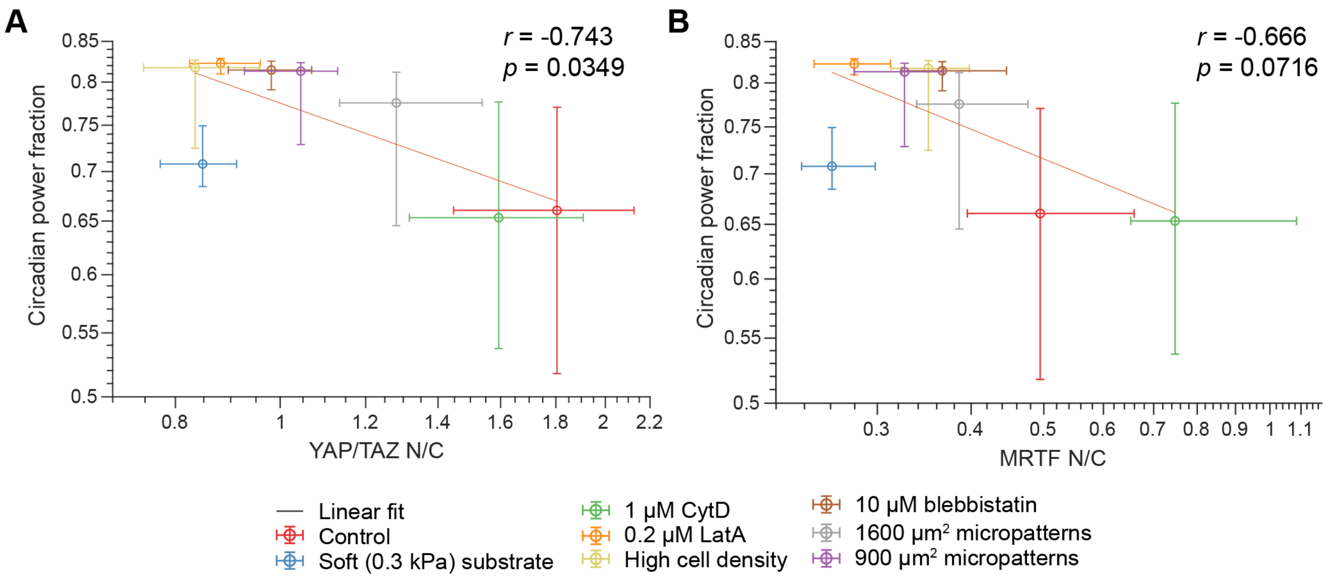

**Figure S4: Altered power fraction / transcriptional regulator correlation with increased expression of PER/CRY and BMAL1 on soft substrates.** Increasing the baseline expression of PER/CRY and BMAL1 for cells on a soft substrate changes the correlations between circadian power fraction and YAP/TAZ N/C (A) or the MRTF N/C (B). In this case, we added additional baseline expression terms to Equations 9 and 10, each $0.0175$ hr$^{-1}$. Pearson's correlation coefficient (between median YAP/TAZ or MRTF N/C and median circadian power fraction, both on the log scale), $r$, and the $p$-value associated with the null hypothesis, $r = 0$, are included on each graph. Data points denote median and error bars denote the range from the 40th to the 60th percentile for 200 model cells. To generate distinguishable colors for each condition, we used the linspecer tool in MATLAB (Lansey 2023).

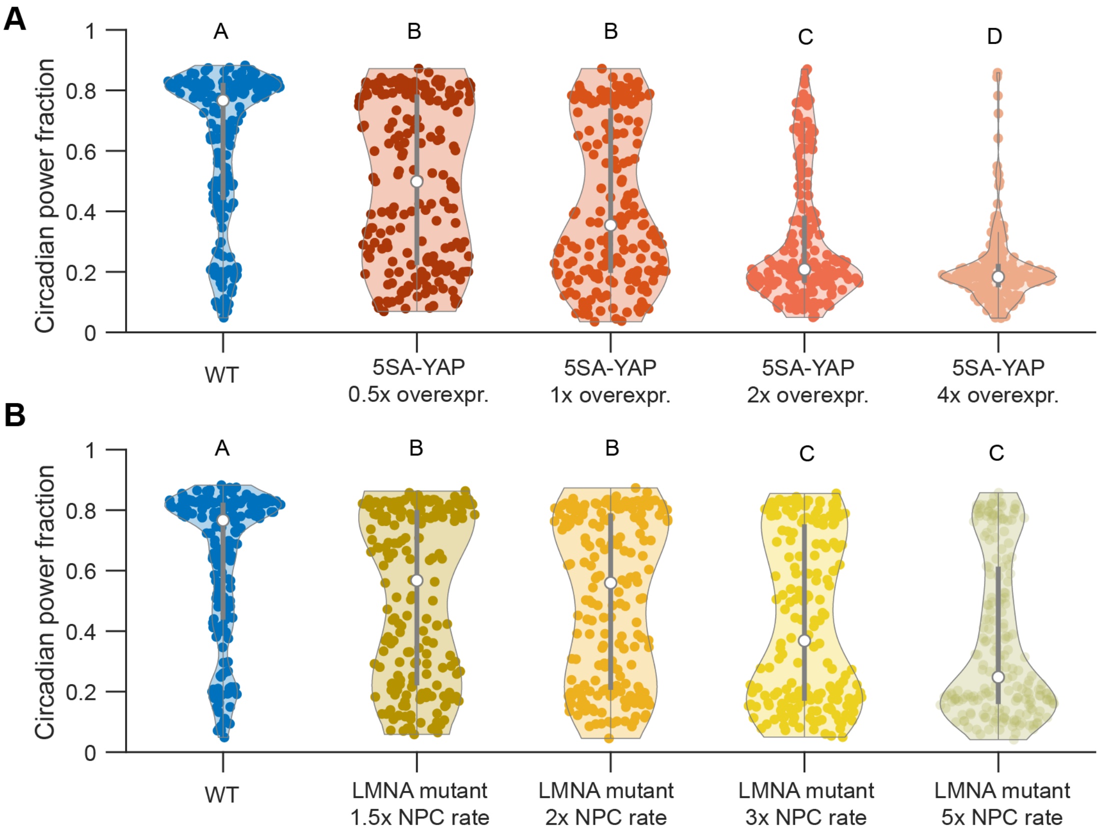

**Figure S5: Effects of alternative mutation parameters on model predictions.** Circadian power fractions plotted for cells with different overexpression levels of YAP mutant (A) and different changes in NPC opening rate induced by LMNA mutation (B). 5SA-YAP overexpression is controlled by $K_{overexpress}$ and NPC opening rate is altered by changing $K_{fNPC}$ (see Tables S2, S3 and S5). Compact letter display is used to denote statistical significance, where groups sharing a letter are statistically similar according to ANOVA followed by Tukey's *post hoc* test with a significance threshold of $p = 0.05$.

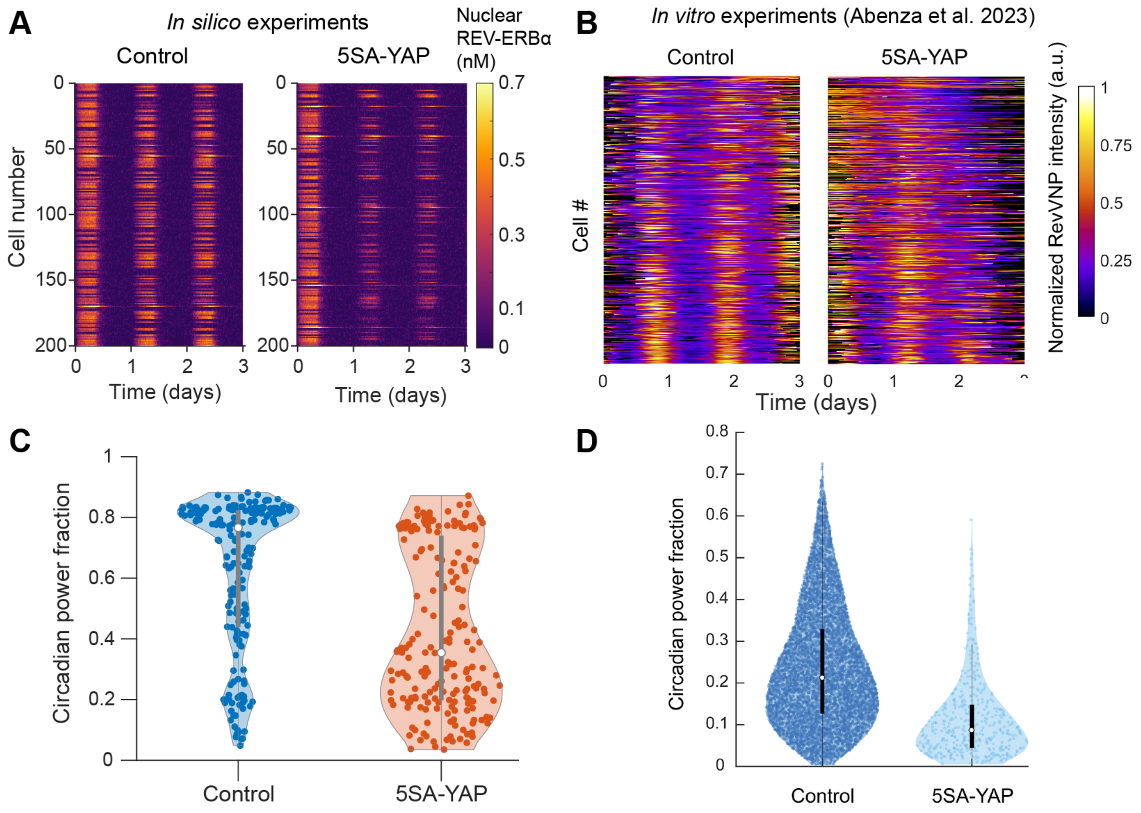

**Figure S6:** Comparison of circadian oscillations for 5SA-YAP mutant in our study vs. Abenza et al. 2023 A) Kymographs depicting the dynamics of BMAL1 in *in silico* cell populations (wild type or 5SA-YAP mutants) on a 30 kPa substrate. Oscillations are substantially weaker for the population of mutant cells. B) Kymographs depicting the REV-VNP dynamics in *in vitro* cell populations (wild type or 5SA-YAP mutants) on a 30 kPa substrate. Oscillations are short-lived and/or inconsistent in the mutant population. C) Circadian power fraction for cells in each simulated population from panel A. 5SA-YAP mutants show a substantial reduction in the power fraction compared to control cells. D) Circadian power fraction for each population in the experiments shown in panel B. Circadian power fraction is much lower for 5SA-YAP mutant cells compared to control, similar to simulated populations. Panels B and D were published in *Journal of Cell Biology*; these plots were taken unaltered from Figure 4 in Abenza et al. 2023 with the publisher's permission.
